# Emission rates of species-specific volatiles change across communities of *Clarkia* species: Evidence for character displacement in floral scent

**DOI:** 10.1101/2020.04.02.022004

**Authors:** Katherine E. Eisen, Monica A. Geber, Robert A. Raguso

**Affiliations:** Department of Ecology and Evolutionary Biology, Cornell University, Ithaca NY, 14853, USA; Department of Neurobiology and Behavior, Cornell University, Ithaca NY, 14853, USA

**Keywords:** allopatry, sympatry, multimodal floral traits, context-dependent selection

## Abstract

A current frontier of character displacement research is to determine if displacement occurs via multiple phenotypic pathways and varies across communities with different species compositions. Here, we conducted the first test for context-dependent character displacement in multimodal floral signals by analyzing variation in floral scent in a system that exhibits character displacement in flower size, and that has multiple types of sympatric communities. In a greenhouse common garden experiment, we measured quantitative variation in volatile emission rates of the progeny of two species of *Clarkia* from replicated communities that contain one, two, or four *Clarkia* species. The first two axes of a constrained correspondence analysis, which explained 24 percent of the total variation in floral scent, separated the species and community types, respectively. Of the 23 compounds that were significantly correlated with these axes, nine showed patterns consistent with character displacement. Two compounds produced primarily by *C. unguiculata* and two compounds produced primarily by *C. cylindrica* were emitted in higher amounts in sympatry. Character displacement in some volatiles varied across sympatric communities and occurred in parallel with displacement in flower size, demonstrating that this evolutionary process can be context-dependent and may occur through multiple pathways.

## Introduction

Interspecific interactions have long been hypothesized to have significant effects on patterns of biodiversity (Darwin 1859; Lack 1945; Schluter 2000; Grant and Grant 2008). One such evolutionary process is character displacement, which leads to a pattern of differences in species’ trait values in sympatric communities relative to allopatric communities (Brown and Wilson 1956; Germain et al. 2017). While character displacement has been studied and debated for over sixty years (Stuart and Losos 2013), there are two key gaps in our understanding of this process. First, outside of a small number of classic systems (e.g., anoles, sticklebacks, Darwin’s finches), few studies have examined the potential for character displacement in more than one type of trait (reviewed in Stuart and Losos 2013). Examining variation in multiple types of traits increases our ability to detect non-repeatable character displacement, which may occur through different phenotypic pathways across communities (Losos 2011; Germain et al. 2017), and determine when species interactions lead to shifts in correlated or independently-evolving traits. Second, while most studies of character displacement have focused on pairwise interactions (but see Lemmon and Lemmon 2010; Miller et al. 2014*a*; Grant 2017; Roth-Monzon et al. 2020), many species exist in complex ecological communities, where interactions with multiple species could include indirect and higher-order interactions (Mayfield and Stouffer 2017; TerHorst et al. 2018; Roth-Monzon et al. 2020). Testing for character displacement across sympatric communities that vary in species composition or richness (Eisen and Geber 2018; Roth-Monzon et al. 2020) can advance our understanding of the evolutionary consequences of direct and indirect interactions (Walsh 2013; TerHorst et al. 2015).

Among co-occurring flowering plants, pollinators often represent a shared resource that is critical for reproduction (Waser et al. 1996; Ollerton et al. 2011), and there is a growing body of evidence for character displacement in plants mediated by interactions between co-occurring species that share pollinators (reviewed in Beans 2014; Eisen and Geber 2018). Presently, there are two critical gaps in our understanding of this process. First, studies to date have examined character displacement in floral morphology, color, and phenology (reviewed in Beans 2014; Eisen and Geber 2018), which reflects a general bias towards visual traits in pollination (Raguso 2008*a*). Nonetheless, olfactory and reward traits are critical to successful pollination in many systems (Schiestl 2010, 2015; Raguso 2014) but have yet to be integrated in to the study of character displacement. Second, character displacement in floral traits is likely to occur via multiple phenotypic pathways or changes in trait combinations across communities (Losos 2011; Germain et al. 2017). Nonetheless, most studies to date have addressed character displacement in one type of floral trait (e.g., color or morphology), not the multi-modal bouquet flowers typically present (Leonard et al. 2011).

The emission of floral scent—volatile organic compounds including monoterpenes, sesquiterpenes, and aromatic compounds (Knudsen et al. 2006)—is a complex trait, in that individual plants can exhibit qualitative variation in the blend of volatiles and quantitative variation in their emission rates (Raguso 2008*b*). Because scent can be produced not only from petals but also from reproductive floral structures, scent may be correlated with or unrelated to variation in flower size (Effmert et al. 2006; Valdivia and Niemeyer 2006; Burdon et al. 2015; Martin et al. 2017), which could lead to multi-modal character displacement in some systems. In addition, species may vary in common volatiles that are produced by a small number of biosynthetic pathways (Dudareva and Pichersky 2006), and in species-specific volatiles that provide ‘private channels’ for communication with specialist pollinators (Raguso 2008*b*; Soler et al. 2010). Insights from three areas of floral scent research suggest that floral scent could undergo character displacement. First, floral scent exhibits substantial intraspecific variation across populations in multiple systems, including cacti (Schlumpberger and Raguso 2008), cycads (Suinyuy et al. 2012), saxifrages (Friberg et al. 2019), and orchids (Gross et al. 2016; Chapurlat et al. 2018). These patterns suggest that floral scent may be relatively evolutionarily labile. As a result, scent could evolve in response to geographic variation in selection (Gross et al. 2016), which could lead to variation in character displacement across different communities (Germain et al. 2017; Eisen and Geber 2018). Second, floral scent can be a target of pollinator-mediated selection (Parachnowitsch et al. 2012; Chapurlat et al. 2019), which indicates that floral scent could evolve in response to interactions between co-occurring plant species that share pollinators. Third, differences in floral scent can mediate reproductive isolation between co-occurring species (Waelti et al. 2008; Bischoff et al. 2014; Peakall and Whitehead 2014) and explain variation in the structure of plant-pollinator networks (Junker et al. 2010; Larue et al. 2016; Kantsa et al. 2018, 2019). As such, floral scent may determine how pollinators are partitioned among co-occurring plant species.

In this study, we test for variation in multimodal character displacement across sympatric communities that contain different numbers of co-occurring species. Specifically, we assess the potential for character displacement in the floral scent of two co-occurring species of California native annuals in the genus *Clarkia* (Onagraceae). These species, *C. unguiculata* and *C. cylindrica*, co-occur more frequently than expected by chance in the southern foothills of the Sierra Nevada (Kern County, CA, USA). Where they co-occur, these species have converged in flowering time and diverged in flower size (Eisen and Geber 2018). This pattern of divergence in flower size provides an opportunity to test if character displacement occurs on multimodal floral signals. We conducted a greenhouse common garden experiment to measure quantitative variation in volatile emission rates of the progeny of plants from natural communities that contain one, two, or four *Clarkia* species. By eliminating variable environmental effects on trait values, the common garden enabled us to compare phenotypes across different, replicated community types and test for significant interactions between species and community types on floral volatile emission rates. These data were used to address three questions regarding the potential for and nature of character displacement in floral scent:

(1): Is variation in volatile emissions across species and community types consistent with character displacement?
(2): Do patterns of character displacement vary across types of sympatric communities? (e.g., two-species vs. four-species communities)
(3): Do multi-modal signals (e.g., floral scent and flower size) jointly undergo character displacement?

## Methods

### Study system

Species in the genus *Clarkia* (Onagraceae) often co-occur and share pollinators, which are primarily solitary bee pollinators that specialize on the genus (Lewis 1953; MacSwain et al. 1973; Singh 2014). Across the genus, species exhibit intra- and interspecific variation in multiple types of floral traits, including flowering time (Lewis 1961; Jonas and Geber 1999; Moeller 2004; Gould et al. 2014; Singh 2014), floral orientation (Lewis 1961), petal coloration (Lewis and Lewis 1955), flower size (Eisen and Geber 2018), and floral scent (Miller et al. 2014*b*). In the Southern Sierra Nevada (Kern River Canyon, Kern County, CA), *C. unguiculata* Lindley and *C. cylindrica* ssp. *clavicarpa* W. Davis co-occur more frequently than expected by chance (Eisen and Geber 2018). These species are primarily outcrossing because flowers are protandrous and herkogamous, and while they share pollinators (Singh 2014), they are not known to hybridize in the field (MacSwain et al. 1973). The petal area of *C. cylindrica* exhibits divergent character displacement (an increase in petal area) relative to *C. unguiculata* in communities that contain two and four species of *Clarkia* (Eisen and Geber 2018).

### Common garden source community selection

Our common garden contained three replicates of each of four unique types of source communities: *C. cylindrica* alone, *C. unguiculata* alone, *C. cylindrica* and *C. unguiculata* together, and these two species with the two other outcrossing *Clarkia* species (*C. speciosa* and *C. xantiana*) that occur in the Kern River Canyon (for community locations, see Table S1). In other words, individuals of each species (*C. cylindrica* and *C. unguiculata*) were grown from seeds sourced from three single-species communities, three two-species communities, and three four-species communities. Community types thus vary in how many species are present in the community. Seeds from both species were collected at all communities in 2017. Three or more fruits per plant were collected from 50 haphazardly chosen plants of each species at each community. The seeds from one fruit from each of 20 plants per community and species were combined to ensure that plants in the common garden represented a sufficient range of any possible plant-level variation at each community.

### Plant germination and growth

Because of the large number of community x species combinations present in the common garden, seeds were started in five batches in September-November 2017. All community x species combinations were included in each batch of plants. To break dormancy, seeds were placed on moist filter paper in a petri dish, wrapped in parafilm, and stratified at 5°C for five to seven days and then held at 23°C for five to seven days before planting. Germinants were transplanted into 656 ml^3^ Cone-tainers (Stuewe & Sons, Tangent, Oregon, USA) filled with Lambert soil mix. The pots’ positions on the greenhouse bench were randomized. Plants were exposed to supplemental light (16 h days) and maintained at 23-25°C during the day and 19-21°C at night. Plants were watered twice a week on average and received on average 30-40 mL of water per week in weeks 1-3 post transplanting, 70-80 mL per week in weeks 4-6, and 100 mL per week in weeks 7-10. Each pot initially contained two germinants; pots were thinned after four weeks to contain one plant. At this time, six prills of Osmocote® Smart-Release® Plant Food Flower & Vegetable 14-14-14 fertilizer (Scotts Miracle-Gro Company, Marysville, OH) were applied to the soil surface in each pot.

### Qualitative scent analysis

To inform our quantitative sampling protocols, we conducted two types of qualitative analyses using Solid Phase Micro Extraction (SPME) fibers (Supelco, Inc., (Sigma-Aldrich), Bellefonte, PA) (Appendix 1). First, to determine if the presence of additional flowers changed the composition of the volatile profile (i.e., due to threshold dosage effects), we compared the profiles of samples with three versus six cut flowers from the same plant. We recovered significantly more monoterpenoid and sesquiterpenoid compounds in samples with six flowers (Appendix 1). Given this result, we adjusted our quantitative headspace sampling protocol to include a minimum of six open flowers per plant (see below). Second, to determine where volatiles are produced in these flowers, we compared the volatile profiles of dissected petals from six flowers versus those of the remaining tissues of the same six flowers. We found that petals generally contained fewer volatiles than the non-petal floral tissues (Appendix 1), which may influence the relationship between flower size and floral scent (see Discussion).

### Quantitative scent analysis

Floral volatile samples were collected using the dynamic headspace adsorption technique between November 13, 2017, and February 5, 2018. All collections were made under natural lighting conditions in a well-aerated glassed-in corridor adjacent to the greenhouse where the plants were grown. We used an AIRCARE hygrometer (Essick Air Products, Little Rock, AR, USA) to record the minimum and maximum temperature and percent humidity during sampling; the average minimums were 17 °C and 23 percent humidity, while the average maximums were 25 °C and 41 percent humidity. Floral samples were obtained from fifteen plants per community per species (*N*_*total*_ = 270), and one vegetative control sample was collected per community per species (*N*_*total*_ = 18).

We used 16 oz PET water bottles to enclose stems for headspace sampling. Water bottles were washed with odorless soap, dried, and baked in a clean drying oven for 15-20 mins at 80 °C each morning before sampling began. Samples were collected using PAS-500 Micro Air Sampler pumps (Spectrex, Inc., Redwood City, CA) connected to traps that contained 0.0100 g of Tenax 80/100 adsorbent (Alltech Associates, Inc. (W.H. Grace), Deerfield, IL, USA). The flow rate of the pump was set to 200 mL/min. Scent was collected for 6 hours, from 900 to 1500, as this corresponds to the period of greatest pollinator activity in natural communities. For floral samples, 6-13 flowers were enclosed per plant (average: 7.4 flowers), and the number of flowers enclosed was recorded for each plant sample. Vegetative controls were obtained from plants that had not begun to flower but had formed buds. During each sampling day, one ambient control sample was collected in an empty PET bottle.

Immediately following the end of the headspace collection period, the traps were removed from the pumps and eluted with 300 μL of GC-MS quality hexane (Burdick & Jackson GC2; Honeywell International, Inc. USA). Samples were then concentrated to 50 μL with a flow of gaseous N_2_, and spiked with 23 ng of toluene (5 mL of a 0.03% solution in hexane) as an internal standard in preparation for analysis with gas chromatography - mass spectrometry (GC-MS; see below). Samples were stored at −20C and labeled with a community-neutral identifier code based on the date of sampling (e.g., December 15-1, December 15-2, etc.) to facilitate analysis that would be blind to the species and community type (see Becklin et al. 2011).

### Scent analysis via GC-MS

Both solvent-eluted and solvent-free (SPME) volatile samples were analyzed using a GC17A gas chromatograph coupled with a QP5000 quadrupole mass spectrometer (Shimadzu Scientific Instruments, Inc., Kyoto, Japan). One μL aliquots of the solvent eluted samples were injected (splitless mode) at 240C onto a polar GC column (EC Wax, 30m long, 0.25 mm internal diameter, 0.25μ film thickness; BGB Analytik). The GC oven program (40C to 240C, increasing at 20C per minute, with a 2-minute hold at the maximum temperature) was optimized to minimize run length (for over 300 samples) while allowing for peak resolution to baseline. Electrical ionization mass spectra were generated under 70eV conditions (scanning range 40-350 m/z), and resulting mass spectra were compared with those of MS libraries (Wiley, NIST, Adams) using Shimadzu GCMSolutions software. Kovats retention indices (KI) were prepared for each compound by running a blend of n-alkanes (C7-C30) under the same chromatographic conditions and optimized method. Volatile compounds were identified via 1) direct comparison of retention time and mass spectra with those of authentic standards, 2) comparison with the KI of the best MS library fit for the unknown with published KI values from the plant volatile literature (NIST WebBook; https://webbook.nist.gov/), or 3) in the absence of standards or published KI values, the mass spectral data (ion fragment table) were listed for unknown compounds in reverse order of abundance, starting with the base peak (set to 100%).

#### Extraction and processing of quantitative data

Peak areas were integrated manually using Shimadzu GCMSolutions software. After excluding compounds that were present in one or two samples out of 270, our quantitative dataset contained 54 compounds (Table S2). To exclude experimental artifacts from individual plants’ profiles, we compared the profile of each sample to the profile of the ambient control that was collected on the same day. If a compound appeared in both a floral sample and the relevant ambient control, we only retained this peak in the floral sample if the floral sample peak area was at minimum five times larger than the peak area of the ambient control. This value was selected to be highly conservative in terms of the compounds that we retained in samples where those compounds were also present in the control. Similarly, to exclude compounds emitted by vegetation, we compared each floral sample to the vegetative control collected from the sample population and applied the same 5x threshold to any overlapping compounds. As such, some compounds were retained in the dataset but excluded from particular samples where their emission rates were similar to the relevant ambient or vegetative controls.

Emission rates were normalized by dividing total ion current (TIC) peak areas by that of the internal standard (Svensson et al. 2005), then were calculated algebraically using response factors generated using external standard dose-response curves generated from log- and semi-log dilutions of the primary floral volatiles identified in these analyses ((*E*)-β-ocimene, α-pinene, β-caryophyllene, benzyl alcohol, and methyl salicylate).

To relate emission rates to floral masses, we measured the fresh and dry masses of twenty flowers (ten male-phase and ten female-phase flowers) per community and species (Table S1). Flowers were selected haphazardly from between four and eight plants per community and species. Each plant contributed a maximum of five flowers to the 20 total flowers per community and species. Fresh masses were recorded immediately after removing the flower from the plant. Flowers were dried for 24 hrs at 50 °C before dry masses were recorded. We present analyses of floral scent emission rates that were standardized by the number of open flowers that contributed to a sample multiplied by the average fresh mass of a flower from that community and species, which gives the μg scent per g fresh floral mass per hr. Analyzing the data using emission rates that were standardized by the number of open flowers that contributed to a sample (μg scent per flower per hr) yielded highly similar results (results not shown).

### Additional common garden to test for wounding artifacts

We observed differences across species and community types in compounds that are generally considered “green leafy volatiles” (abbreviated as GLVs) and are associated with plant wounding (Visser and Ave 1978; Scala et al. 2013) (see Results). To determine if these patterns resulted from artifacts of the experimental sampling process, we conducted an additional common garden experiment to test for differences in the emission rates of GLVs between wounded and non-wounded plants (see Appendix 2). B Wounding elevated the emission rates of GLVs (e.g., (*Z*)-3-hexen-1-ol, (*Z*)-3-hexenyl acetate), but the emission rates of these compounds during the 2018 main experiment were more similar to the 2019 non-wounded control plants than to the 2019 wounded plants (see Appendix 2). As such, observed emission rates of these compounds are unlikely to be an experimental artifact and we retain the GLVs as floral compounds in our analysis.

### Multivariate statistical analyses and dimensionality reduction

We used multivariate and univariate methods to test for significant interactions between species and community type, which provides evidence for differences in species’ emission rates of volatiles across communities (character displacement). All analyses were performed in R (R Core Team 2018). To partition the observed variance in the emission rate of all compounds across the species and community types, we performed a Permutational Multivariate Analysis of Variance (PERMANOVA) using a Bray-Curtis distance matrix using the adonis function from the vegan package (Oksanen et al. 2018). We specified our replicate communities nested within community type as strata in the function, which is equivalent to a random effect. To find the compounds that distinguish the community types and species, we performed a Canonical Analysis of Principal Coordinates with a Bray-Curtis distance dissimilarity index using the capscale function from the vegan package (Oksanen et al. 2018).

### Univariate statistical analyses

We analyzed variation in compound classes and in specific compounds using general linear mixed-effects models, which were performed using the lme4 package (Bates et al. 2015). Models were assessed to ensure normally distributed residuals with homogenous variance. These models all contained community type, species, and their interaction as fixed effects, and community nested within community type was included as a random effect. The significance of fixed effects in linear mixed models was assessed using the ANOVA function in the lmerTest package ver. 2.0-29 (Kuznetsova et al. 2015) to perform type III F tests using the Kenward-Roger approximation for the denominator degrees of freedom. When ANOVAs returned significant F values, we used Tukey’s honest significant difference tests to determine which group means were significantly different using the emmeans function with the pairwise option in the emmeans package (Lenth 2019). These tests were performed with the ‘type = “response”’ option such that intervals were back-transformed from the log and square-root scales. Contrasts for models with log-transformed response variables are presented on the log-odds scale, such that ratios greater than one indicate larger emission rates in *C. cylindrica* and ratios less than one indicate larger emission rates in *C. unguiculata*.

We performed two types of univariate analyses. First, we tested for differences in total scent emission and the emission of certain types of compounds across the species and community types using linear mixed-effects models as described above. The compound classes we analyzed were monoterpenoids, sesquiterpenoids, GLVs, and aromatics (Table S2). To ensure that our models had normally distributed residuals with homogenous variance, total scent, monoterpenoid, and aromatic emission rates were square-root transformed and GLVs and sesquiterpenoid emission rates were log-transformed.

Second, we performed univariate analyses on compounds that were correlated with one or both of the first two CAP axes. Specifically, 23 compounds were correlated with one or both of the first two CAP axes (Table S3). These compounds had significant Pearson correlation coefficients with one or both axes at *P* < 0.01 after applying a false discovery rate correction for 54 tests; all significant correlations were greater than 0.15 or less than −0.15. Most compounds were either square-root or log-transformed to improve the normality of model residuals (Table S3). To test for a significant interaction between species and community type, we ran ANOVAs on these models as described above. We applied a false discovery rate correction for 23 tests on the *P* values associated with these species by community type interactions. We then tested for differences between species and community types using the emmeans function as described above. All datasets and analysis scripts are available on github (https://github.com/kate-eisen/clarkia-scent).

## Results

### Diversity of volatile organic compounds

There were 54 volatile organic compounds present in four or more samples: 22 monoterpenoids, 18 sesquiterpenes and C_15_ derivatives, five GLVs and nine aromatic or nitrogenous compounds (Table S2). Thirty-eight of the 54 compounds were found in more than five samples of both species. Of the remaining 16 compounds, 11 compounds were completely or nearly unique to *C. unguiculata* (present in five or fewer *C. cylindrica* samples), and five compounds were completely or nearly unique to *C. cylindrica* (present in five or fewer *C. unguiculata* samples). The average number of compounds detected in a sample (mean ± 1 SE) was 13.8 ± 0.4 for *C. cylindrica* and 14.0 ± 0.5 for *C. unguiculata*.

### Multivariate analyses

The PERMANOVA on the scent compounds revealed main effects of community type (R^2^ = 0.03, *P* < 0.001), species (R^2^ = 0.22; *P* < 0.001), and an interaction between the two (R^2^ = 0.04, *P* < 0.001).

The Canonical Analysis of Principal Coordinates indicated that a subset of all compounds helped to define variation among the six species x community type combinations in our study (two focal species × three community types per species). The constrained portion of the variance was 24% of the total variance (our independent variables explained 24 % of the total variation in the data). CAP axis 1 explained 20 % of the total variation in scent and 77 % of the constrained variance. In general, *C. cylindrica* individuals had negative values on CAP axis 1, while *C. unguiculata* individuals had positive values (Figure 1, Table S4). CAP axis 1 was strongly positively correlated with two monoterpenoids and an aromatic compound, and strongly negatively correlated with sesquiterpenoids (Table 1, Table S3). CAP axis 2 primarily separated *C. unguiculata* individuals from the three different community types (Table S4). This axis explained 4 % of the total variation in scent, and 14 % of the constrained variance. It was strongly positively correlated with two monoterpenoids and an aromatic compound, and strongly negatively correlated with two GLVs and a monoterpenoid (Table 1, Table S3).

**Figure 1.**
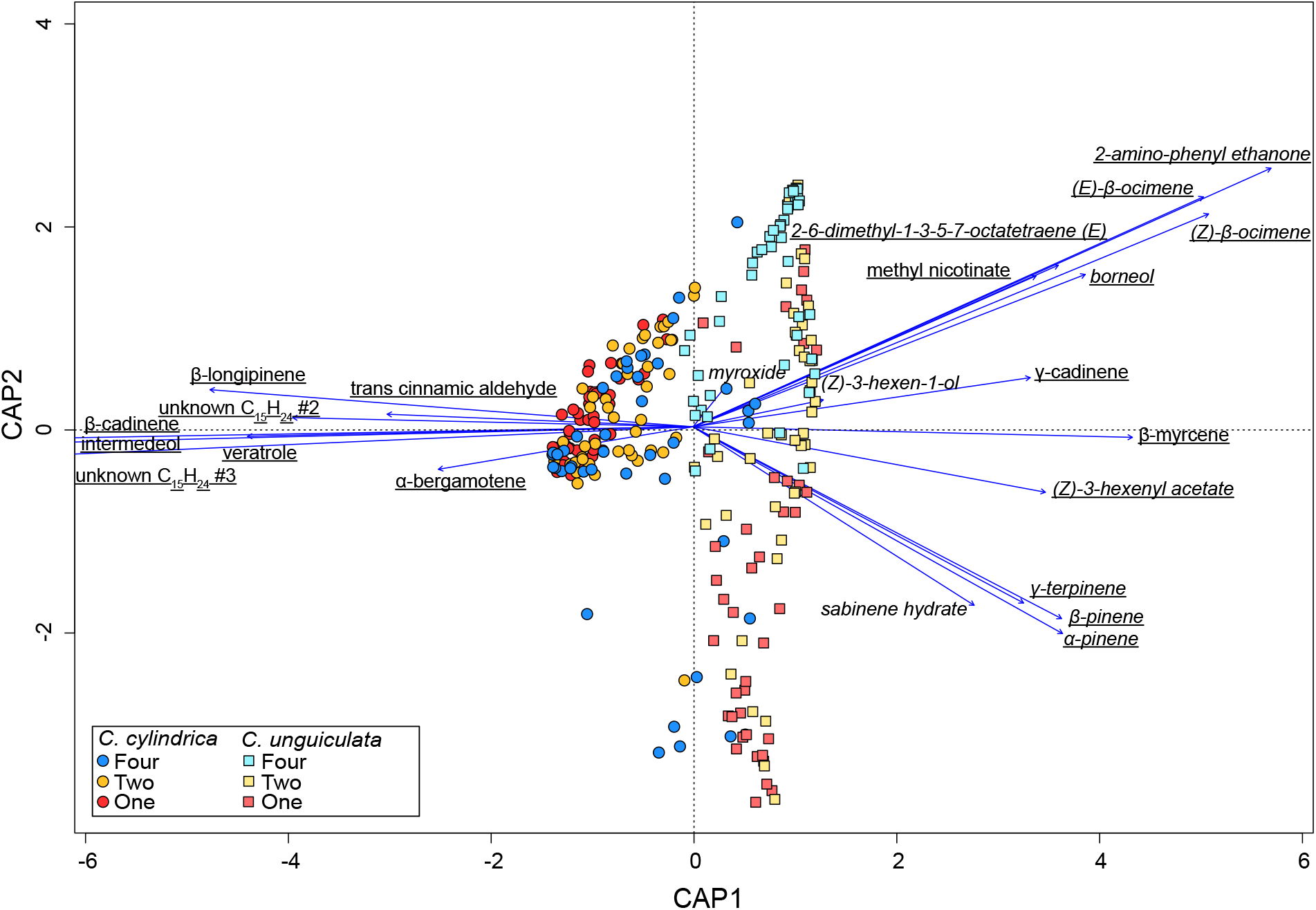
Canonical analysis of principle coordinates separating volatile emissions from the two species and three community types. The different species and community type combinations are indicated by color and symbol type. The loadings are shown with arrows for compounds with the largest positive and negative correlations (*r*) with CAP axis 1 (underlined compounds), and for compounds with the largest positive and negative correlations (*r*) with CAP axis 2 (italicized compounds). CAP axis 1 strongly separates the species, as *C. cylindrica* have lower values and *C. unguiculata* have higher values. CAP axis 2 primarily differentiates the community types of *C. unguiculata*, as individuals from one-species communities had generally low values (centroid= −1.212), individuals from two-species communities had intermediate values (centroid: −0.061), and individuals from four-species communities had high values (centroid: 1.330).

**Table 1.** The three compounds with the strongest positive and negative correlations with the first two CAP axes. For CAP1, nine additional compounds (two “green leafy volatiles,” five monoterpenoids, one sesquiterpenoid, and one aromatic compound) were significantly positively correlated with this axis, and three additional sesquiterpenoids and two aromatic compounds exhibited significant negative correlations (*r* > 0.15; Table S3). For CAP2, two additional monoterpenoids and one aromatic compound were also positively correlated with this axis, and three additional monoterpenes exhibited significant negative correlations (*r* > 0.15; Table S3). A false discovery rate correction for conducting 54 tests was applied to all *P* values (see Methods).

| Axis and compound | $r$ | $P$ |
| --- | --- | --- |
| CAP1 |  |  |
| (*Z*)-β-ocimene | 0.528 | $1.51 \times 10^{-19}$ |
| (*E*)-β-ocimene | 0.526 | $1.70 \times 10^{-19}$ |
| 2-amino phenyl ethanone | 0.481 | $4.60 \times 10^{-16}$ |
| intermedeol | -0.646 | $1.73 \times 10^{-31}$ |
| β-cadinene | -0.641 | $2.87 \times 10^{-31}$ |
| unknown C<sub>15</sub>H<sub>24</sub> #3 | -0.516 | $1.04 \times 10^{-18}$ |
| CAP2 |  |  |
| (*E*)-β-ocimene | 0.457 | $1.26 \times 10^{-13}$ |
| (*Z*)-β-ocimene | 0.444 | $5.16 \times 10^{-13}$ |
| (*E*)-2-6-dimethyl-1,3,5,7- octatetraene | 0.329 | $4.12 \times 10^{-7}$ |
| (*Z*)-3-hexenyl acetate | -0.349 | $6.44 \times 10^{-8}$ |
| (*Z*)-3-hexen-1-ol | -0.275 | $3.52 \times 10^{-5}$ |
| sabinene hydrate | -0.275 | $3.52 \times 10^{-5}$ |

### Univariate analyses of total scent, compound classes, and single compounds

Patterns of variation in three compound classes were consistent with character displacement: sesquiterpenes (*F*_2,117.84_ = 8.749, *P* = 0.0003), GLVs (*F*_2,78.42_ = 17.330, *P* < 0.0001), and aromatics (*F*_2,89.51_ = 4.720, *P* = 0.0113). The interaction in sesquiterpene emissions was driven by a significantly larger difference between the species in one-species communities relative to two-species communities (Figure 2; Table S5). For the GLVs, *Clarkia unguiculata* produced more than *C. cylindrica* at one-species communities and two-species communities; the interaction was driven by both species producing equivalent amounts of these compounds at four-species communities (Figure 2; Table S5). The interaction in aromatics emissions was driven by greater production by *C. cylindrica* at two-species communities (Figure 2; Table S5), such that the difference between the species at two-species communities was significantly larger than the differences between the species at one- and four-species communities (Figure 2; Table S5). In particular, this pattern was driven by the emission of large amounts of benzyl alcohol by *C. cylindrica* in two-species communities (results not shown).

**Figure 2.**
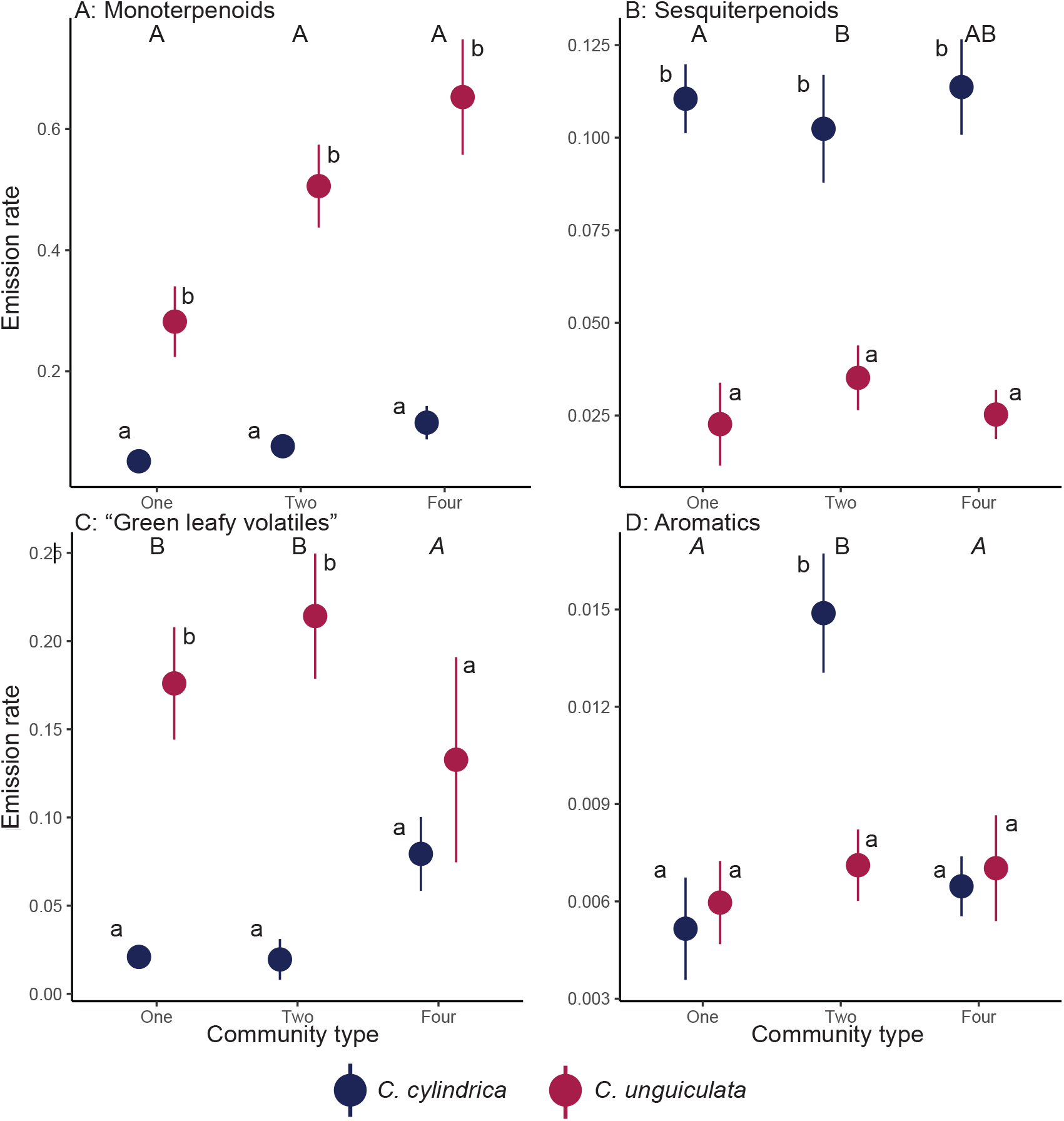
Emission rates (raw values; μg scent/g fresh floral mass/hour) of monoterpenoids (A), sesquiterpenoids (B), “green leafy volatiles” (C), and aromatic compounds (D) by *C. cylindrica* (navy blue) and *C. unguiculata* (dark pink). Lowercase letters above each point indicate whether emission rates differed between the species within that community type; the letter b indicates the species with the higher emission rate. Uppercase letters above each set of points at a community type indicate whether that difference is the same or different from the differences at other community types; differences with the letter A in italics are not significantly different from zero, and differences with the letters B or C are larger than differences with the letter A. Emission rates of monoterpenoids were higher in *C. unguiculata* (A). *Clarkia cylindrica* produced more sesquiterpenes than *C. unguiculata* at all community types (B), but the difference between the species was significantly larger at one-species communities than at two-species communities. *Clarkia unguiculata* produced more “green leafy volatiles” than *C. cylindrica* at one-species communities and at two-species communities, but emission rates at four-species communities were equivalent (C). *Clarkia cylindrica* had substantially higher emission rates of aromatic compounds at two-species communities (D). Note the differences in scale for the y-axes across all panels.

We used the results of the Canonical Analysis of Principal Coordinates to determine the compounds that we analyzed individually. Of the 23 compounds that were correlated with one or both of the first two CAP axes (see Methods), nine compounds had significant community type by species interactions in univariate models (Table S6). Two of these compounds, 2-amino phenyl ethanone, and methyl nicotinate, are primarily produced by *C. unguiculata*, and emission rates were higher at both types of sympatric communities (Figure 3 A&B, Table S5). Two additional compounds, (*E*)-cinnamic aldehyde and veratrole, are primarily or exclusively produced by *C. cylindrica*, and emission rates were higher at both types of sympatric communities (Figure 3 G & H, Table S5). The remaining five compounds with significant interactions, α-pinene, β-pinene, sabinene hydrate, γ-terpinene, and (*Z*)-3-hexenyl acetate, are primarily produced by *C. unguiculata* and had lower emission rates in four-species communities relative to one- and two-species communities (Figure 3, Table S5).

**Figure 3.**
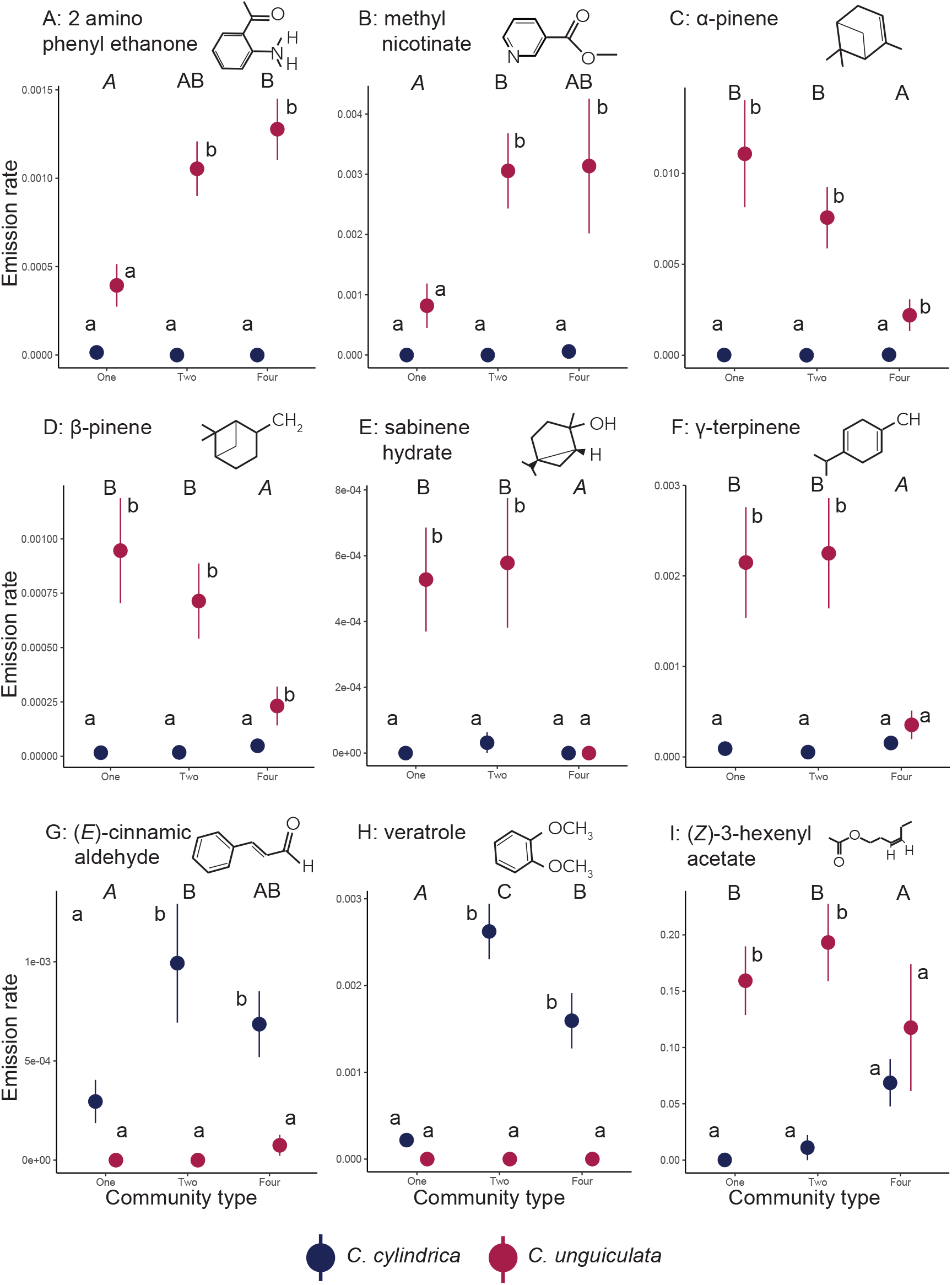
Emission rates (raw values; μg scent/g fresh floral mass/hour) of the nine compounds that were significantly correlated with one or both of the first two CAP axes and also had a significant species x community type interaction. Lowercase letters above each point indicate whether emission rates differed between the species within that community type; the letter b indicates the species with the higher emission rate. Uppercase letters above each set of points at a community type indicate whether that difference is the same or different from the differences at other community types; differences with the letter A in italics are not significantly different from zero, and differences with the letters B or C are larger than differences with the letter A. 2-amino-phenyl ethenone (A) and methyl nicotinate (B) were primarily emitted by *C. unguiculata* (dark pink) and increased in two- and four-species communities. α-pinene (C), β-pinene (D), sabinene hydrate (E), and γ-terpinene (F) were also primarily emitted by *C. unguiculata* and decreased in four-species communities. (*E*)-cinnamic aldehyde (G) and veratrole (H) were exclusively emitted by *C. cylindrica* (navy blue) and the emission rates of these compounds increased in two- and four-species communities. (*Z*)-3-hexenyl acetate (I) was emitted at higher rates by *C. unguiculata* at one- and two-species communities, and by *C. cylindrica* at four-species communities. Note the differences in scale for the y-axes across all panels.

## Discussion

By measuring floral scent variation across communities that contain different numbers of species in a system that exhibits character displacement in flower size, we conducted the first test for context-dependent multi-modal character displacement in floral traits. These species exhibit pronounced differences in their floral scent profiles, with more subtle but significant differences across the community types. In an analysis of all of the volatile organic compounds emitted by the two species, the significant interaction between species and community type was driven by compounds that were primarily or exclusively emitted by only one species—two aromatic compounds and four monoterpenoids emitted by *C. unguiculata*, and two aromatic compounds emitted by *C. cylindrica*. These patterns were consistent across sympatric communities for *C. cylindrica* but not for *C. unguiculata*. In addition, our investigation of the potential for multi-modal character displacement revealed that changes in floral scent were associated with changes in flower size in *C. cylindrica* but not in *C. unguiculata.* Here we discuss the potential drivers and ecological implications of these patterns.

### Character displacement driven by changes in species-specific volatiles

Because floral scent is a complex trait, character displacement could occur through several pathways, including both qualitative or quantitative changes in compounds that are either shared across the species or unique to one species. In this study, we observed patterns consistent with character displacement in compounds that were generally emitted by only one of the focal species. These types of changes could be linked to increases in plant-pollinator specialization in multi-species communities. Specifically, an increase in species-specific volatile emissions may increase a pollinator’s ability to differentiate between two co-occurring plant species, which could increase pollinator constancy and decrease heterospecific pollen transfer among species that share pollinators (Waser 1986; Sargent and Ackerly 2008). Divergence in flower color has been demonstrated to reduce inconstant foraging in multiple systems (Levin 1985; Hopkins and Rausher 2012; Muchhala et al. 2014; Norton et al. 2015). The compounds that exhibited patterns consistent with character displacement in *Clarkia* were benzenoid aromatics (both species) and monoterpenoids (*C. unguiculata*). Among plants that are pollinated by food-seeking bees, scent profiles are commonly dominated by benzenoids, terpenoids, or a mixture of the two types of compounds (Dobson 2006). In both observational and experimental studies, benzenoids have been associated with visitation from apid and halictid bees (Theis 2006; Andrews et al. 2007; Kantsa et al. 2019), such that the increases in benzenoid emission rates in *Clarkia* could result in greater attraction of these pollinator species. In particular, because only *C. unguiculata* receives upwards of five percent of all pollinator visits from apid bees (*Apis mellifera*, *Xylocopa tabaniformis*, *Bombus* sp.; Singh 2014), the increases in benzenoid emissions could reflect greater pollinator specialization in sympatric communities.

### Context dependency of character displacement

Because indirect interactions can modify evolutionary trajectories (Benkman 2013; Walsh 2013; TerHorst et al. 2015), we tested for variation in character displacement in two types of sympatric communities: two-species communities that contain the focal species of this study, and four-species communities that contain the focal species plus two additional congeners that flower later in the summer (Moeller 2004). We found that *C. cylindrica* exhibited similar patterns across both types of sympatric communities, while patterns for *C. unguiculata* across the community types varied by compound class (monoterpenoids and aromatics). In general, this variation in the patterns observed for our two focal species points to the potential for character displacement to be context-dependent (Eisen and Geber 2018; Roth-Monzon et al. 2020) and to occur via different phenotypic pathways across communities (Germain et al. 2017). In particular, our results suggest that changes in the volatile profile of *C. cylindrica* may result primarily from interactions with *C. unguiculata*, which occurs in both types of sympatric communities. For *C. cylindrica*, indirect interactions with the later-flowering *Clarkia* species in the four-species communities may not affect the evolution of floral scent.

In contrast, *C. unguiculata* had greater emission rates of two aromatic compounds at both types of sympatric communities, but lower emission rates of four monoterpenoids only at the four-species sympatric communities. Similar patterns of intermediate or less displacement were observed across different multispecies communities of freshwater fish (Roth-Monzon et al. 2020), which suggests that evolution in these communities likely occurs in response to multiple species interactions. Because *C. unguiculata* is the earliest *Clarkia* species to flower in the region (Moeller 2004; Singh 2014), its higher total scent emission (see Figure 2) may serve to attract scarce pollinators at the beginning of the flowering season (Filella et al. 2013). However, the observed decrease in monoterpenoid emissions in the four-species communities suggests that *C. unguiculata* may invest less in pollinator attraction if, like other species of *Clarkia* (Moeller 2004), it experiences facilitation in these communities.

### Multi-modal character displacement: synergy of changes in floral size & scent

Because pollinators often exhibit responses to combinations of visual and olfactory traits (Leonard et al. 2011), we conducted the first test for character displacement in multi-modal floral signals (Figure 4). Using estimates of volatile emission rates that were standardized by floral fresh mass, we found that changes in the floral scent of *C. unguiculata* were not associated with changes in flower size, while increases in the emission of floral scent of *C. cylindrica* were related to increases in flower size. We hypothesize that this pattern results from differences in the floral parts that produce these compounds (Effmert et al. 2006). Our floral dissections suggest that the character displacement compounds in *C. cylindrica* are produced in both the petals and the reproductive parts, which is consistent with a correlation between flower size and floral scent. In contrast, the character displacement compounds in *C. unguiculata* were present more often in the reproductive parts relative to the petals (Appendix 1), which is consistent with a change in floral volatiles that was independent of a change in flower size. The differences in these patterns highlight that the complexity of floral scent can extend beyond the quantitative and qualitative composition of a scent bouquet to include spatial variation in the emission of volatiles across tissue types (Friberg et al. 2013; Burdon et al. 2015; Martin et al. 2017).

**Figure 4.**
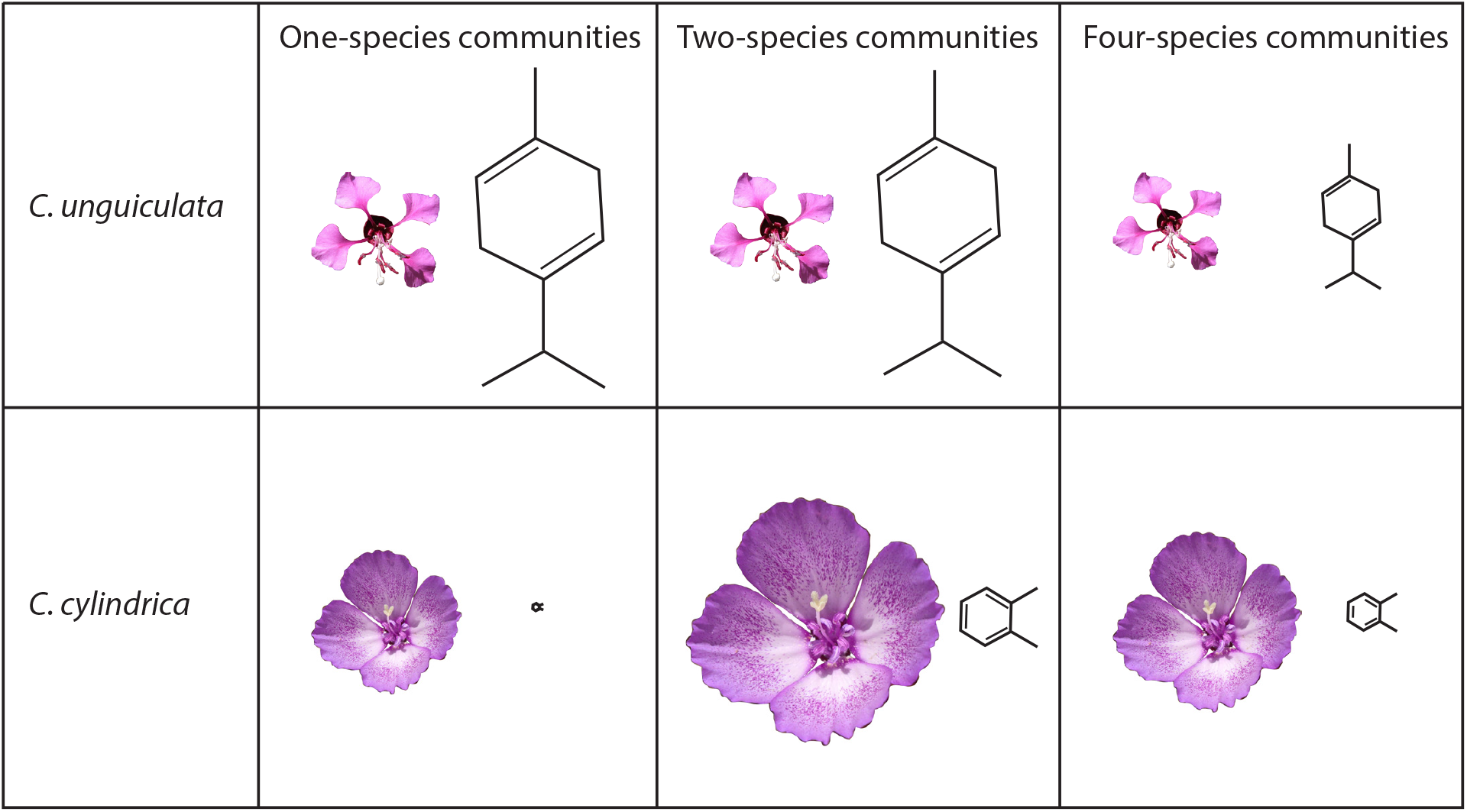
Schematic showing the relative changes in flower size (based on measurements of petal area in Eisen & Geber 2018) and the species-specific floral scent compounds that showed patterns consistent with character displacement (*C. unguiculata*: 2-amino phenyl ethenone, α-pinene, β-pinene, sabinene hydrate, γ-terpinene, and methyl nicotinate; *C. cylindrica*: (*E*)- cinnamic aldehyde and veratrole). Drawings of flowers and chemical compounds (molecules that are representative of the suites of compounds that responded in each species) are scaled proportionally both between the species and across the community types. Flower size of *C. unguiculata* is similar across community types and is slightly smaller than the flower size of *C. cylindrica* at one-species communities. Floral scent emission rates of *C. unguiculata* are similar at one- and two-species communities, and emission rates at four-species communities are about 0.45 times emission rates at one species-communities (a decrease in scent emission in four-species communities). Emission rates of floral scent in *C. unguiculata* are one or two orders of magnitude higher than emission rates in *C. cylindrica.* Flower size of *C. cylindrica* at two-species communities is 1.7 times larger than flower size at one-species communities, and flower size at four-species communities is 1.25 times larger than at one-species communities. Floral scent emission rates of *C. cylindrica* are 7.2 times larger at two-species communities relative to one-species communities, and 4.6 times larger at four-species communities relative to one-species communities.

These differences in the floral sources of the volatiles that change across the community types may signify differences in their functions. For *C. cylindrica*, the increases in both size and volatile emissions in both petals and reproductive parts may serve to increase overall pollinator attraction in sympatry. Increases in flower size or scent emission have been linked to increased pollinator attraction and plant reproductive success in multiple insect-pollinated systems (Conner and Rush 1996; Miyake and Yafuso 2003; Majetic et al. 2009; Sandring and Ågren 2009; Parachnowitsch et al. 2012), although most studies have not tested for concurrent changes in both traits (but see Parachnowitsch et al. 2012). For *C. unguiculata*, scent emission in the reproductive tissues may serve to cue pollinators to the precise location of the reproductive parts (Dötterl and Jürgens 2005; Burdon et al. 2015). Because the solitary bees that specialize on *Clarkia* forage for pollen (MacSwain et al. 1973), volatiles emitted in the reproductive tissues also indicate the location of the primary rewards for this species. After becoming attracted to a flower, bees can use pollen odors, which are often a distinct subset of the floral bouquet (Jürgens and Dötterl 2004; Effmert et al. 2006), to orient more specifically to the source of pollen (Dobson et al. 1996, 1999). Here, the decrease in floral volatiles that are putatively produced in the reproductive organs in four-species communities suggests that *C. unguiculata* may invest less not only in pollinator attraction as described above, but more specifically in provisioning pollinators with pollen where the community of congeners may facilitate joint pollinator attraction or maintenance (Moeller 2004). This hypothesis could be tested via additional analysis of the pollen volatiles in *C. unguiculata*, and with pollinator behavior assays (see below).

### Future directions

This study yielded a pattern of trait variation that is consistent with character displacement, but additional work is needed to rule out alternative hypotheses (Schluter and McPhail 1992). In particular, it is critical to determine if this variation in scent has functional consequences for pollinator behavior. Given that the volatiles that mediate pollinator behavior are often a subset of all volatiles emitted by a plant (reviewed in Junker and Blüthgen 2010), pollinators may not respond to the specific changes observed in floral scent profiles across community types. Experimental assays of pollinator behavior can be used to determine if these shifts in volatiles affect pollinator attraction or constancy, or if they are non-functional. The potential effects of variation in *C. unguiculata* volatiles on honey bees and bumblebees could be tested in a controlled environment (e.g., Burger et al. 2012; Peter and Johnson 2014). However, a comprehensive assessment of the functionality of floral scent variation in *Clarkia* would need to be field-based, as lab experiments with the solitary bees that pollinate both species are not tractable.

More broadly, the results of this study highlight the need to continue to integrate chemical phenotypes into the study of floral trait evolution (Leonard et al. 2011; Junker and Parachnowitsch 2015). In combination with visual traits, floral scent can affect species interactions at multiple scales, from specifying highly specialized interactions (e.g., Peakall and Whitehead 2014; Whitehead et al. 2015) to contributing to the structure of complex plant-pollinator interaction networks (Kantsa et al. 2018, 2019). In this study, by testing for character displacement in multiple trait modalities across communities that contain different numbers of interacting species, we have generated new insights into the context-dependency of character displacement, which may occur through multiple pathways. Moving forward, systems that exhibit variation in both floral scent and species interactions across communities (e.g., Friberg et al. 2019) provide opportunities to study the interplay between complex trait evolution and species interactions, which can generate insight into the repeatability of evolutionary change across variable ecological communities.

## Supporting information

All supplemental material (appendices and tables)

## Acknowledgements

We thank Geoff Broadhead and Amy Wruck for assistance in the lab and greenhouse. We thank Erika Mudrak from the Cornell Statistical Consulting Center and Diane Campbell for advice on statistical analyses. Anurag Agrawal provided insightful comments on drafts of the manuscript. This work was funded by a Graduate Research Fellowship (DGE-1650441) from the US National Science Foundation, and a Cornell Atkinson Center for Sustainability Sustainable Biodiversity Fund grant to KEE, by NSF DEB-1754299 to MAG, and by NSF DEB-1342792 to RAR.

## References

Andrews, E. S., N. Theis, and L. S. Adler. 2007. Pollinator and herbivore attraction to *Cucurbita* floral volatiles. Journal of Chemical Ecology 33:1682–1691.

Bates, D., M. Mächler, B. M. Bolker, and S. C. Walker. 2015. Fitting linear mixed-effects models using lme4. Journal of Statistical Software 67:1–48.

Beans, C. M. 2014. The case for character displacement in plants. Ecology and Evolution 4:852–65.

Becklin, K. M., G. Gamez, B. Uelk, R. A. Raguso, and C. Galen. 2011. Soil fungal effects on floral signals, rewards, and aboveground interactions in an alpine pollination web. American Journal of Botany 98:1299–1308.

Benkman, C. W. 2013. Biotic interaction strength and the intensity of selection. Ecology Letters 16:1054–1060.

Bischoff, M., A. Jürgens, and D. R. Campbell. 2014. Floral scent in natural hybrids of *Ipomopsis* (Polemoniaceae) and their parental species. Annals of Botany 113:533–544.

Brown, W. L., and E. O. Wilson. 1956. Character displacement. Systematic Zoology 5:49–64.

Burdon, R. C. F., R. A. Raguso, A. Kessler, and A. L. Parachnowitsch. 2015. Spatiotemporal floral scent variation of *Penstemon digitalis*. Journal of Chemical Ecology 41:641–650.

Burger, H., S. Dötterl, C. M. Häberlein, S. Schulz, and M. Ayasse. 2012. An arthropod deterrent attracts specialised bees to their host plants. Oecologia 168:727–736.

Chapurlat, E., J. Agren, J. Anderson, M. Friberg, and N. Sletvold. 2019. Conflicting selection on floral sent emission in the orchid *Gymnadenia conopsea*. New Phytologist 222:2009–2022.

Chapurlat, E., J. Anderson, J. Ågren, M. Friberg, and N. Sletvold. 2018. Diel pattern of floral scent emission matches the relative importance of diurnal and nocturnal pollinators in populations of *Gymnadenia conopsea*. Annals of Botany 121:711–721.

Conner, J. K., and S. Rush. 1996. Effects of flower size and number on pollinator visitation to wild radish, Raphanus raphanistrum. Oecologia 105:509–516.

Darwin, C. 1859. The origin of species. John Murray, London.

Dobson, H. E. M. 2006. Relationship between floral fragrance composition and type of pollinator. Pages 147–198 in E. Pichersky and N. Dudareva, eds. Biology of Floral Scent. Chapman & Hall.

Dobson, H. E. M., E. M. Danielson, and I. D. Van Wesep. 1999. Pollen odor chemicals as modulators of bumble bee foraging on *Rosa rugosa* Thunb. (Rosaceae). Plant Species Biology 14:153–166.

Dobson, H. E. M., I. Groth, and G. Bergström. 1996. Pollen advertisement: chemical contrasts between whole-flower and pollen odors. American Journal of Botany 83:877–885.

Dötterl, S., and A. Jürgens. 2005. Spatial fragrance patterns in flowers of *Silene latifolia*: Lilac compounds as olfactory nectar guides? Plant Systematics and Evolution 255:99–109.

Dudareva, N., and E. Pichersky. 2006. Floral scent metabolic pathways: Their regulation and evolution. Pages 55–78 in E. Pichersky and N. Dudareva, eds. Biology of Floral Scent. Chapman & Hall.

Effmert, U., D. Buss, D. Rohrbeck, and B. Piechulla. 2006. Localization of the synthesis and emission of scent compounds within the flower. Pages 105–124 in N. Dudareva and E. Pichersky, eds. Biology of Floral Scent. CRC Press, Boca Raton, FL.

Eisen, K. E., and M. A. Geber. 2018. Ecological sorting and character displacement contribute to the structure of communities of *Clarkia* species. Journal of Evolutionary Biology 31:1440–1458.

Filella, I., C. Primante, J. Llusià, A. M. Martín González, R. Seco, G. Farré-Armengol, A. Rodrigo, et al. 2013. Floral advertisement scent in a changing plant-pollinators market. Scientific Reports 3:1–6.

Friberg, M., C. Schwind, P. R. Guimarães, R. A. Raguso, and J. N. Thompson. 2019. Extreme diversification of floral volatiles within and among species of *Lithophragma* (Saxifragaceae). Proceedings of the National Academy of Sciences 201809007.

Friberg, M., C. Schwind, R. A. Raguso, and J. N. Thompson. 2013. Extreme divergence in floral scent among woodland star species (*Lithophragma* spp.) pollinated by floral parasites. Annals of Botany 111:539–550.

Germain, R. M., J. L. Williams, D. Schluter, and A. L. Angert. 2017. Moving character displacement beyond characters using contemporary coexistence theory. Trends in Ecology & Evolution 33:74–84.

Gould, B., D. A. Moeller, V. M. Eckhart, P. Tiffin, E. Fabio, and M. A. Geber. 2014. Local adaptation and range boundary formation in response to complex environmental gradients across the geographical range of *Clarkia xantiana* ssp. *xantiana*. Journal of Ecology 102:95–107.

Grant, P. R. 2017. Ecology and Evolution of Darwin’s Finches. Princeton University Press, Princeton, New Jersey, USA.

Grant, P. R., and B. R. Grant. 2008. How and why species multiply: the radiation of Darwin’s finches. Princeton University Press, Princeton, NJ.

Gross, K., M. Sun, and F. P. Schiestl. 2016. Why do floral perfumes become different? Region-specific selection on floral scent in a terrestrial orchid. PLoS ONE 11:e0147975.

Hopkins, R., and M. D. Rausher. 2012. Pollinator-mediated selection on flower color allele drives reinforcement. Science 335:1090–1092.

Jonas, C. S., and M. A. Geber. 1999. Variation among populations of *Clarkia unguiculata* (Onagraceae) along altitudinal and latitudinal gradients. American Journal of Botany 86:333–343.

Junker, R. R., and N. Blüthgen. 2010. Floral scents repel facultative flower visitors, but attract obligate ones. Annals of Botany 105:777–782.

Junker, R. R., N. Höcherl, and N. Blüthgen. 2010. Responses to olfactory signals reflect network structure of flower-visitor interactions. Journal of Animal Ecology 79:818–823.

Junker, R. R., and A. L. Parachnowitsch. 2015. Working towards a holistic view on flower traits-how floral scents mediate plant-animal interactions in concert with other floral characters. Journal of the Indian Institute of Science 95:43–67.

Jürgens, A., and S. Dötterl. 2004. Chemical composition of anther volatiles in Ranunculaceae: Genera-specific profiles in *Anemone*, *Aquilegia*, *Caltha*, *Pulsatilla*, *Ranunculus*, and *Trollius* species. American Journal of Botany 91:1969–1980.

Kantsa, A., R. A. Raguso, A. G. Dyer, J. M. Olesen, T. Tscheulin, and T. Petanidou. 2018. Disentangling the role of floral sensory stimuli in pollination networks. Nature Communications 9:1–25.

Kantsa, A., R. A. Raguso, T. Lekkas, O. I. Kalantzi, and T. Petanidou. 2019. Floral volatiles and visitors: A meta-network of associations in a natural community. Journal of Ecology 107:2574–2586.

Knudsen, J. T., R. Eriksson, J. Gershenzon, and B. Ståhl. 2006. Diversity and distribution of floral scent. The Botanical Review 72:1–120.

Kuznetsova, A., P. Brockhoff, and R. Christensen. 2015. Package “lmerTest.” R package version.

Lack, D. 1945. The Galapagos finches (*Geospizinae*): a study in variation. California Academy of Sciences.

Larue, A. A. C., R. A. Raguso, and R. R. Junker. 2016. Experimental manipulation of floral scent bouquets restructures flower-visitor interactions in the field. Journal of Animal Ecology 85:396–408.

Lemmon, E. M., and A. R. Lemmon. 2010. Reinforcement in chorus frogs: Lifetime fitness estimates including intrinsic natural selection and sexual selection against hybrids. Evolution 64:1748–1761.

Lenth, R. 2019. emmeans: Estimated Marginal Means, aka Least-Squares Means. R package version 1.3.3.

Leonard, A. S., A. Dornhaus, and D. R. Papaj. 2011. Forget-me-not: Complex floral displays, inter-signal interactions, and pollinator cognition. Current Zoology 57:215–224.

Levin, D. A. 1985. Reproductive character displacement in *Phlox*. Evolution 39:1275–1281.

Lewis, H. 1953. The mechanism of evolution in the genus *Clarkia*. Evolution 7:1–20.

Lewis, H. 1961. Experimental sympatric populations of *Clarkia*. The American Naturalist 95:155–168.

Lewis, H., and M. E. Lewis. 1955. The genus *Clarkia*. University of California Publications in Botany 20:241–392.

Losos, J. B. 2011. Convergence, adaptation, and constraint. Evolution 65:1827–1840.

MacSwain, J., P. H. Raven, and R. Thorp. 1973. Comparative behavior of bees and Onagraceae. IV. *Clarkia* bees of the western United States. University of California Publications in Entomology 70:1–80.

Majetic, C. J., R. A. Raguso, and T. L. Ashman. 2009. The sweet smell of success: Floral scent affects pollinator attraction and seed fitness in *Hesperis matronalis*. Functional Ecology 23:480–487.

Martin, K. R., M. Moré, J. Hipólito, S. Charlemagne, B. O. Schlumpberger, and R. A. Raguso. 2017. Spatial and temporal variation in volatile composition suggests olfactory division of labor within the trap flowers of *Aristolochia gigantea*. Flora: Morphology, Distribution, Functional Ecology of Plants 232:153–168.

Mayfield, M. M., and D. B. Stouffer. 2017. Higher-order interactions capture unexplained complexity in diverse communities. Nature Ecology & Evolution 1:0062.

Miller, T. E., E. R. Moran, and C. P. TerHorst. 2014a. Rethinking niche evolution: Experiments with natural communities of Protozoa in pitcher plants. The American Naturalist 184:277–283.

Miller, T. J., R. A. Raguso, and K. M. Kay. 2014b. Novel adaptation to hawkmoth pollinators in *Clarkia* reduces efficiency, not attraction of diurnal visitors. Annals of botany 113:317–29.

Miyake, T., and M. Yafuso. 2003. Floral scents affect reproductive success in fly-pollinated *Alocasia odora* (Araceae). American Journal of Botany 90:370–376.

Moeller, D. A. 2004. Facilitative interactions among plants via shared pollinators. Ecology 85:3289–3301.

Muchhala, N., S. Johnsen, and S. Smith. 2014. Competition for hummingbird pollination shapes flower color variation in Andean Solanaceae. Evolution 68:2275–2286.

Norton, N. A., M. T. R. Fernando, C. R. Herlihy, and J. W. Busch. 2015. Reproductive character displacement shapes a spatially structured petal color polymorphism in *Leavenworthia stylosa*. Evolution 69:1191–1207.

Oksanen, J., F. G. Blanchet, M. Friendly, R. Kindt, P. Legendre, D. McGlinn, P. R. Minchin, et al. 2018. vegan: Community Ecology Package. R package version 2.5-2.

Ollerton, J., R. Winfree, and S. Tarrant. 2011. How many flowering plants are pollinated by animals? Oikos 120:321–326.

Parachnowitsch, A. L., R. A. Raguso, and A. Kessler. 2012. Phenotypic selection to increase floral scent emission, but not flower size or colour in bee-pollinated *Penstemon digitalis*. New Phytologist 195:667–675.

Peakall, R., and M. R. Whitehead. 2014. Floral odour chemistry defines species boundaries and underpins strong reproductive isolation in sexually deceptive orchids. Annals of Botany 113:341–355.

Peter, C. I., and S. D. Johnson. 2014. A pollinator shift explains floral divergence in an orchid species complex in South Africa. Annals of Botany 113:277–288.

R Core Team. 2018. R: A language and environment for statistical computing. R Foundation for Statistical Computing, Vienna, Austria.

Raguso, R. A. 2008a. Start making scents: The challenge of integrating chemistry into pollination ecology. Entomologia Experimentalis et Applicata 128:196–207.

Raguso, R. A. 2008b. Wake up and smell the roses: The ecology and evolution of floral scent. Annual Review of Ecology, Evolution, and Systematics 39:549–569.

Raguso, R. A. 2014. A wrapped bouquet: the untapped potential of floral chemistry. Journal of Chemical Ecology 40:412–3.

Roth-Monzon, A. J., M. C. Belk, J. J. Zuniga-Vega, and J. B. Johnson. 2020. Beyond pairwise interactions: multispecies character displacement in Mexican freshwater fish communities. American Naturalist doi: 10.1086/708513.

Sandring, S., and J. Ågren. 2009. Pollinator-mediated selection on floral display and flowering time in the perennial herb *Arabidopsis lyrata*. Evolution 63:1292–1300.

Sargent, R. D., and D. D. Ackerly. 2008. Plant-pollinator interactions and the assembly of plant communities. Trends in Ecology & Evolution 23:123–30.

Scala, A., S. Allmann, R. Mirabella, M. A. Haring, and R. C. Schuurink. 2013. Green leaf volatiles: A plant’s multifunctional weapon against herbivores and pathogens. International Journal of Molecular Sciences 14:17781–17811.

Schiestl, F. P. 2010. The evolution of floral scent and insect chemical communication. Ecology Letters 13:643–656.

Schiestl, F. P. 2015. Ecology and evolution of floral volatile-mediated information transfer in plants. New Phytologist 206:571–577.

Schlumpberger, B. O., and R. A. Raguso. 2008. Geographic variation in floral scent of *Echinopsis ancistrophora* (Cactaceae); evidence for constraints on hawkmoth attraction. Oikos 117:801–814.

Schluter, D. 2000. The ecology of adaptive radiation. Oxford University Press.

Schluter, D., and J. D. McPhail. 1992. Ecological character displacement and speciation in sticklebacks. The American Naturalist 140:85–108.

Singh, I. 2014. *Pollination interaction networks between* Clarkia *(Onagraceae) species and their pollinators in the Southern Sierra Nevada, California*. Cornell University.

Soler, C., M. Proffit, C. Chen, and M. Hossaert-McKey. 2010. Private channels in plant-pollinator mutualisms. Plant Signaling and Behavior 5:893–895.

Stuart, Y. E., and J. B. Losos. 2013. Ecological character displacement: Glass half full or half empty? Trends in Ecology and Evolution 28:402–408.

Suinyuy, T. N., J. S. Donaldson, and S. D. Johnson. 2012. Geographical variation in cone volatile composition among populations of the African cycad *Encephalartos villosus*. Biological Journal of the Linnean Society 106:514–527.

Svensson, G. P., M. O. Hickman, S. Bartram, W. Boland, O. Pellmyr, and R. A. Raguso. 2005. Chemistry and geographic variation of floral scent in *Yucca filamentosa* (Agavaceae). American Journal of Botany 92:1624–1631.

TerHorst, C. P., J. A. Lau, I. A. Cooper, K. R. Keller, R. J. La Rosa, A. M. Royer, E. H. Schultheis, et al. 2015. Quantifying non-additive selection caused by indirect ecological effects. Ecology 96:2360–2369.

TerHorst, C. P., P. C. Zee, K. D. Heath, T. E. Miller, A. I. Pastore, S. Patel, S. J. Schreiber, et al. 2018. Evolution in a community context: Trait responses to multiple species interactions. The American Naturalist 191:368–380.

Theis, N. 2006. Fragrance of Canada thistle (*Cirsium arvense*) attracts both floral herbivores and pollinators. Journal of Chemical Ecology 32:917–927.

Valdivia, C. E., and H. M. Niemeyer. 2006. Do pollinators simultaneously select for inflorescence size and amount of floral scents? An experimental assessment on *Escallonia myrtoidea*. Austral Ecology 31:897–903.

Visser, J., and D. Ave. 1978. General green leaf volatiles in the olfactory orientation of the Colorado beetle, *Lerptinotarsa decemlineata*. Entomologia Experimentalis et Applicata 24:738–749.

Waelti, M. O., J. K. Muhlemann, A. Widmer, and F. P. Schiestl. 2008. Floral odour and reproductive isolation in two species of *Silene*. Journal of Evolutionary Biology 21:111–121.

Walsh, M. R. 2013. The evolutionary consequences of indirect effects. Trends in Ecology and Evolution 28:23–29.

Waser, N. M. 1986. Flower constancy: definition, cause, and measurement. The American Naturalist 127:593–603.

Waser, N. M., L. Chittka, M. V Price, N. M. Williams, and J. Ollerton. 1996. Generalization in pollination systems, and why it matters. Ecology 77:1043–1060.

Whitehead, M. R., C. C. Linde, and R. Peakall. 2015. Pollination by sexual deception promotes outcrossing and mate diversity in self-compatible clonal orchids. Journal of Evolutionary Biology 28:1526–1541.

