## Supplementary material for "Emission rates of species-specific volatiles change across communities of *Clarkia* species: Evidence for character displacement in floral scent": All supplemental material (appendices and tables)

Appendix 1: Supplemental methods and results from qualitative SPME sampling

*Rationale.* We conducted two types of additional analyses using Solid Phase Micro Extraction (SPME) fibers (Supelco, Inc., (Sigma-Aldrich), Bellefonte, PA). To determine if the presence of additional flowers changed the composition of the volatile profile (i.e. due to threshold effects), we compared the profiles of samples with three versus six cut flowers from the same plant. To determine if volatiles are produced in the petals and/or reproductive parts of these flowers, we compared the volatile profiles of dissected petals from six flowers versus those of the remaining tissues of the same six flowers.

*Methods.* We used the 65 μ "Stabilflex" field-assembly fibers (with both divinylbenzene and polydimethylsiloxane in the adsorbent matrix) because of their proven versatility in trapping different biosynthetic classes of volatile compounds (Goodrich & Raguso, 2009). Both types of collections were performed on plants of both species from one one-species community (*C. cylindrica*: MHG; *C. unguiculata*: CB323) and from one two-species community (SC for both species). Three replicates of each comparison were sampled for each analysis (e.g. 3 samples with 3 flowers and 3 samples with 6 flowers from each community for each species).

Samples were collected in 20 mL glass scintillation vials sealed with Nalophan (PTE) film (Toppits ®, Cofresco Frischhalteprodukte, Minden, Germany). One ambient control vial was sampled during each sampling period. All samples were equilibrated for 60 minutes, and exposed to a SPME fiber for 30 minutes. After the exposure period, fibers were inserted into the GC injection port for thermal desorption. Analysis via GC-MS followed the same method used for the solvent-eluted samples.

Peak areas were integrated manually using Shimadzu GCMSolutions software. We observed 45 compounds across all samples (Table S2), and compounds were identified using the same protocols as the solvent-eluted samples (see Methods). To exclude experimental artifacts, each sample was compared to the concurrently collected ambient control. No ambient control samples contained the compounds detected in the floral samples.

*Statistical analysis.* To determine if the presence of additional flowers changed the composition of the volatile profile (i.e. due to threshold effects), we compared the number of compounds observed in the three flower samples versus the six flower samples. Specifically, we calculated the total number of compounds in each sample, and the number of monoterpenoids, sesquiterpenoids, aromatics, and "green leafy volatiles" (see Table S2). To test for differences in these count data, we ran paired Wilcoxon signed rank tests for each compound class.

To determine if volatiles are produced in the petals and/or reproductive parts of these flowers, we compared the presences and absences of all compounds across petal and non-petal samples. We performed a Permutational Multivariate Analysis of Variance (PERMANOVA) using a jaccard distance matrix on the jaccard dissimilarity values between samples using the adonis function from the vegan package in R (Oksanen et al., 2019). We visualized the differences between the petal and non-petal samples using the multivariate dispersion, which shows the average distance to the group centroid.

*Results*. Samples with six flowers contained significantly more compounds than samples with three flowers ($Z$ = -2.4382, $P$ = 0.015). Specifically, the samples with six flowers contained more monoterpenoids ($Z$ = -2.4945, $P$ = 0.013), and sesquiterpenoids ($Z$ = -2.1264, $P$ = 0.033). In the PERMANOVA analysis of samples from petals and non-petals, there was a significant effect of sample type ($R_2$ = 0.1555, $P$ = 0.002). Petal and non-petal samples formed two distinct clusters based on their multivariate dispersion (Figure 1). Non-petal samples contained significantly more compounds than petal samples (*Z* = 3.0618, *P* = 0.0022).

*Conclusions*. The significant increase in monoterpenoids and sesquiterpenoids in the samples with six flowers suggests that increasing the floral tissue in a sample can increase the probability of detecting a fuller complement of compounds, more representative of a blooming inflorescence. As such, we collected quantitative samples from plants with six or more open flowers in our study. The dissected flower tissues were separated in multivariate space, which suggests that scent is differentially produced across types of floral tissue in both species. In particular, the non-petal samples contained more compounds than the petal samples, which suggests that increases in volatile production may not be strongly correlated with increases in petal size in these species.

*References:*


Goodrich, K. R., and R. A. Raguso. 2009. The olfactory component of floral display in *Asimina* and *Deeringothamnus* (Annonaceae). New Phytologist 183:457–469.

Oksanen, J., Blanchet, F. G., Friendly, M., Kindt, R., Legendre, P., McGlinn, D., Minchin, P. R., O'Hara, R. B., Simpson, G. L., Solymos, P., Stevens, M. H. H., Szoecs, E., and H. Wagner. 2019. vegan: Community Ecology Package. R package version 2.5-6. https://CRAN.R-project.org/package=vegan

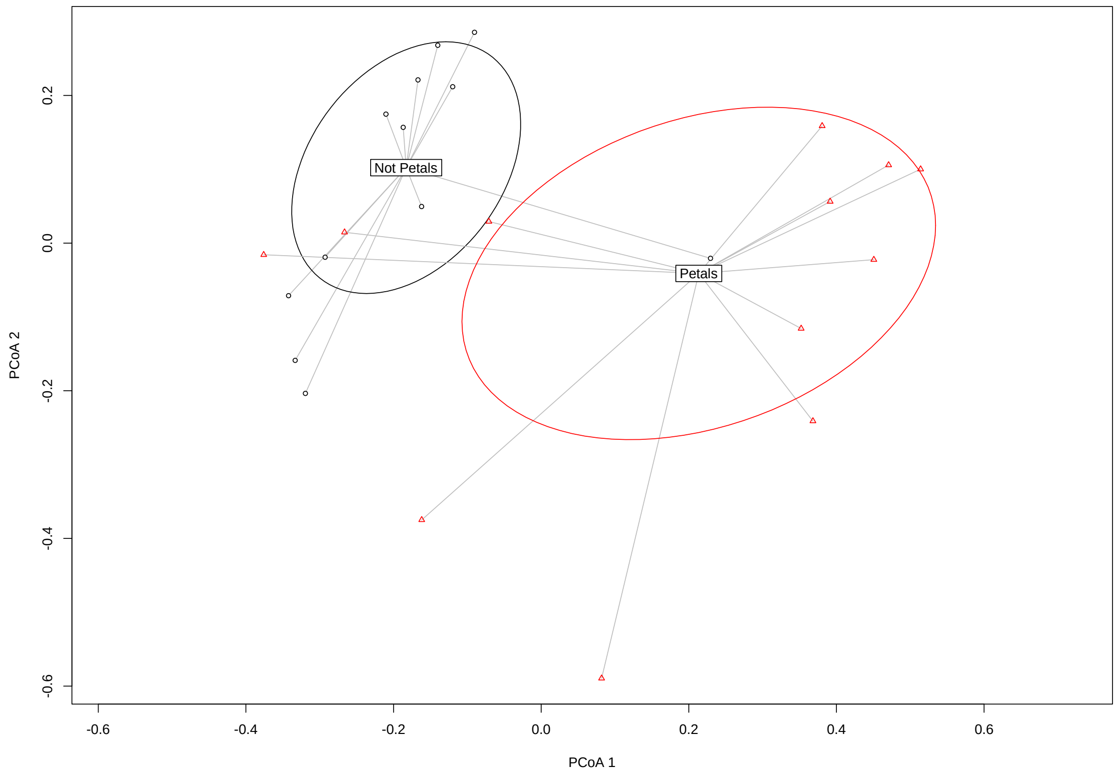


Figure 1. Multivariate dispersion of the petal and non-petal samples, performed using jaccard distances. The ellipses show the group standard deviations, and lines represent the average distance to the group centroid.

Appendix 2: Supplemental methods and results from tests for wounding artifacts

*Rationale*. We observed differences across species and community types in compounds that are generally considered "green leafy volatiles" associated with plant wounding (Visser and Ave 1978; Scala et al. 2013). To determine if these patterns were the product of an experimental artifact, we conducted a second greenhouse common garden experiment to compare the volatiles emitted by wounded and non-wounded plants.

*Plant source community selection*. We grew individuals of both species from seeds from one single-species community per species (*C. cylindrica*: MHG; *C. unguiculata*: GRCO), one two-species community per species (SC for both species), and one four-species community per species (MCK for both species). These communities were specifically selected to represent a range in the emission rates of the "green leafy volatiles" observed in the 2018 common garden.

*Plant germination and growth*. Seeds were stratified and transplanted in two batches in January and February of 2019. All procedures were identical to those used in the 2018 common garden.

*Quantitative scent analysis*. Floral volatile samples were collected using the dynamic headspace adsorption technique between March 21, 2019 and April 6, 2019. All collection protocols were identical to those used in the 2018 common garden. To assess the potential for elevated emission of "green leafy volatiles" due to wounding, each plant was sampled twice. On the first sampling day for a given plant, we sampled the floral bouquet using the same methods as in 2018, to produce a control sample. On the second sampling day for a given plant, we sampled the floral bouquet and inflicted wounds to the plant immediately prior to the collection window. Using scissors, we snipped a piece of each leaf included in the collection chamber for *C. cylindrica*, and we snipped every other leaf included in the collection chamber for *C. unguiculata* because

*C. unguiculata* plants have more, larger leaves. This resulted in a mean ± 1 SE of 0.065 ± 0.005 g of fresh leaf mass removed from *C. cylindrica* and 0.049 ± 0.004 g of fresh leaf mass removed from *C. unguiculata*. The number of flowers included in each sample was recorded. We collected both non-wounded and wounded samples from 10 plants per species per community ($N_{total}$ = 120 samples).

*Scent analysis via GC-FID*. Samples were analyzed using a Shimadzu GC-FID (GC 2014) with an AOC-20i auto injector. One μL aliquots of the solvent eluted samples were injected (splitless mode) at 240C onto a polar GC column (EC Wax, 30m long, 0.25 mm internal diameter, 0.25μ film thickness; BGB Analytik). The GC oven program (a 2-minute hold at 40C, followed by a 14.67C increase per minute to the maximum temperature of 260C, with a 2-minute hold at the maximum temperature) was optimized to minimize run length (for over 120 samples) while allowing for peak resolution to baseline. The "green leafy volatile" compounds of interest ((*Z*)-3-hexen-1-ol and (*Z*)-3-hexenyl acetate) were identified via direct comparison of retention time and mass spectra with those of authentic standards. Peak areas were automatically integrated by Shimadzu Postrun Analysis software.

*Extraction and processing of quantitative data.* Emission rates were calculated as in the 2018 common garden using response factors generated using external standard dose-response curves generated from log- and semi-log dilutions of the compounds of interest. Emission rates were related to floral masses using the floral mass data collected in 2018.

Emission rates from each sample were compared to the ambient control collected on that day, but none of the ambient control samples contained the compounds of interest.

*Statistical analysis*. The data were subset by species and we used paired t-tests (t.test function in R) to compare the wounded vs. non-wounded volatile emission profiles. We then examined the

2018 common garden data that had been cleaned to remove compounds with emission rates less than 5x the emission rates in the ambient control, but that had not been cleaned to remove compounds that occur in the vegetative controls. We analyzed this form of the 2018 data because we did not utilize vegetative controls in our 2019 common garden, such that the 2018 profiles prior to vegetative control screening are directly comparable to the 2019 profiles. We ran two sets of two sample t-tests for each species: 1) we compared the 2018 profiles to the 2019 non-wounded control samples, and 2) we compared the 2018 profiles to the 2019 wounded samples. Due to the variance structure of these data, we ran two sample t tests with unequal variance for (*Z*)-3-hexen-1-ol and with equal variances for (*Z*)-3-hexenyl acetate).

*Expected results*. If our experimental protocol induces a wounding response, we expect that the "green leafy volatile" emission rates will be equivalent across the 2019 wounded and 2019 non-wounded plants. If our experimental protocol does not induce a wounding response, we expect that the "green leafy volatile" emission rates will be higher in the 2019 wounded samples relative to the 2019 non-wounded samples. In addition, we expect that the 2018 emission rates will be roughly equivalent to the 2019 non-wounded emission rates, and lower than the 2019 wounded emission rates.

*Results and Discussion*. For both species and both compounds, emission rates were higher in the 2019 wounded samples relative to the 2019 non-wounded samples (Table 1), which suggests that wounding elevates emission rates of these volatile compounds. For both species and compounds, emission rates did not differ between the 2019 control samples and the 2018 samples (see 2018 – 2019 C comparisons Table 2), which suggests that our 2019 control samples are representative of the emission rates observed in 2018. For *C. unguiculata*, emissions rates of both compounds were lower in the 2018 samples relative to the 2019 wounded samples (see 2018 – 2019 W comparisons Table 2). Comparisons of emission rates for *C. cylindrica* yield similar patterns, although the comparison for (*Z*)-3-hexen-1-ol is non-significant but in the expected direction. Taken together, these results suggest that while wounding elevates the emission rates of these "green leafy volatiles," the emission rates observed in 2018 cannot be attributed to wounding. Rather, these emission rates likely reflect constitutive emission of these compounds by flowering plants, which has been documented in other systems (e.g., Brodmann et al. 2008, 2012).

*References*


Brodmann, J., D. Emer, and M. Ayasse. 2012. Pollinator attraction of the wasp-flower Scrophularia umbrosa (Scrophulariaceae). Plant Biology 14:500–505.

Brodmann, J., R. Twele, W. Francke, G. Hölzler, Q. H. Zhang, and M. Ayasse. 2008. Orchids mimic green-leaf volatiles to attract prey-hunting wasps for pollination. Current Biology 18:740–744.

Scala, A., S. Allmann, R. Mirabella, M. A. Haring, and R. C. Schuurink. 2013. Green leaf volatiles: A plant's multifunctional weapon against herbivores and pathogens. International Journal of Molecular Sciences 14:17781–17811.

Visser, J., and D. Ave. 1978. General green leaf volatiles in the olfactory orientation of the Colorado beetle, Lerptinotarsa decemlineata. Entomologia Experimentalis et Applicata 24:738–749

Table 1. Results of paired t-tests comparing the 2019 wounded and non-wounded plant samples. Ung refers to *C. unguiculata* and Cyl refers to *C. cylindrica*. CI.low and CI.high represent the lower and upper bounds respectively of a 95% confidence interval on the mean difference. Positive mean differences indicate higher emission rates in the wounded samples. Significant tests are bolded, and marginally significant tests are italicized.

| **Species** | **Compound** | ***t*** | **df** | ***P*** | **CI.low** | **CI.high** | **Mean difference** |
|---|---|---|---|---|---|---|---|
| Ung | (*Z*)-3-hexen-1-ol | *1.7066* | *29* | *0.0986* | -0.0020 | 0.0226 | 0.0103 |
| | (*Z*)-3-hexenyl acetate | **2.5461** | **29** | **0.0164** | 0.0264 | 0.2421 | 0.1342 |
| Cyl | (*Z*)-3-hexen-1-ol | **3.7503** | **29** | **0.0008** | 0.0042 | 0.0145 | 0.0094 |
| | (*Z*)-3-hexenyl acetate | **2.9732** | **29** | **0.0059** | 0.0337 | 0.1825 | 0.1081 |

Table 2. Results of two sample t-tests comparing the 2018 samples to the 2019 non-wounded controls (2019 C) and to the 2019 wounded samples (2019 W). Ung refers to *C. unguiculata* and Cyl refers to *C. cylindrica*. CI.low and CI.high represent the lower and upper bounds respectively of a 95% confidence interval on the mean difference. Mean 2019 refers to the mean of the 2019 group included in the specific analysis. Significant negative *t* values indicate that the 2019 group included in the test had a higher emission rate than the 2018 samples. Significant tests are bolded.

| **Species** | **Compound** | **Comparison** | ***t*** | **df** | ***P*** | **CI.low** | **CI.high** | **Mean 2018** | **Mean 2019** |
|---|---|---|---|---|---|---|---|---|---|
| Ung | (*Z*)-3-hexen-1-ol | 2018 – 2019 C | -1.4014 | 32 | 0.1707 | -0.0209 | 0.0039 | 0.0102 | 0.0188 |
| | | 2018 – 2019 W | **-5.0833** | **38** | **< 0.0001** | -0.0263 | -0.0113 | | 0.0290 |
| | (*Z*)-3-hexenyl acetate | 2018 – 2019 C | -0.0258 | 73 | 0.9795 | -0.1157 | 0.1127 | 0.2022 | 0.2036 |
| | | 2018 – 2019 W | **-2.6321** | **73** | **0.0104** | -0.2385 | -0.0321 | | 0.3379 |
| Cyl | (*Z*)-3-hexen-1-ol | 2018 – 2019 C | 0.6783 | 73 | 0.4997 | -0.0058 | 0.0117 | 0.0156 | 0.0126 |
| | | 2018 – 2019 W | -1.4618 | 73 | 0.1481 | -0.0152 | 0.0023 | | 0.0220 |
| | (*Z*)-3-hexenyl acetate | 2018 – 2019 C | -1.3750 | 73 | 0.1733 | -0.1316 | 0.0241 | 0.1633 | 0.2170 |
| | | 2018 – 2019 W | **-3.9411** | **73** | **0.0002** | -0.2437 | -0.0800 | | 0.3251 |

Supplemental Tables

Table S1. Source communities utilized in the common garden. For species composition, *C* = *C. cylindrica*, *U* = *C. unguiculata*, *S* = *C. speciosa*, and *X* = *C. xantiana.* Floral mass data are the average ± 1 SE of 20 flowers (10 male and 10 female) from that community.

| **Community Name** | **Latitude** | **Longitude** | **Species composition** | **Floral wet mass (g)** | **Floral dry mass (g)** |
|---|---|---|---|---|---|
| Site 33 | 35.4657 | -118.7538 | *U* | 0.127595 ± 0.006931 | 0.032200 ± 0.001976 |
| Green Corner | 35.4615 | -118.7627 | *U* | 0.122480 ± 0.007222 | 0.026620 ± 0.001713 |
| Callbox 323 | 35.5534 | -118.6158 | *U* | 0.100425 ± 0.008205 | 0.021675 ± 0.002006 |
| Main Highway Gulley | 35.5834 | -118.5302 | *C* | 0.092420 ± 0.006989 | 0.019645 ± 0.001506 |
| Upper Coyote Gulch | 35.5816 | -118.5216 | *C* | 0.095650 ± 0.005825 | 0.020295 ± 0.001141 |
| Sandy Flats | 35.5809 | -118.5258 | *C* | 0.096650 ± 0.006690 | 0.020040 ± 0.001265 |
| Summer Camp | 35.5294 | -118.6460 | *U* | 0.085860 ± 0.004403 | 0.016955 ± 0.001074 |
| | | | *C* | 0.132430 ± 0.008060 | 0.027980 ± 0.001578 |
| North Pole | 35.5323 | -118.6472 | *U* | 0.092390 ± 0.004091 | 0.019445 ± 0.000811 |
| | | | *C* | 0.087565 ± 0.005921 | 0.019060 ± 0.001180 |
| Divot | 35.4742 | -118.7286 | *U* | 0.089840 ± 0.005520 | 0.020970 ± 0.001344 |
| | | | *C* | 0.148845 ± 0.010543 | 0.029585 ± 0.002052 |
| Democrat | 35.5289 | -118.6266 | *U* | 0.113970 ± 0.004510 | 0.025620 ± 0.001087 |
| | | | *C* | 0.101840 ± 0.005847 | 0.020300 ± 0.001383 |
| | | | *S* | | |
| | | | *X* | | |
| Delonegha East | 35.5464 | -118.6170 | *U* | 0.091820 ± 0.006289 | 0.020935 ± 0.001475 |
| | | | *C* | 0.075940 ± 0.005110 | 0.015205 ± 0.000982 |
| | | | *S* | | |
| | | | *X* | | |
| Mill Creek | 35.5363 | -118.6142 | *U* | 0.085310 ± 0.004255 | 0.018130 ± 0.001303 |
| | | | *C* | 0.101405 ± 0.006916 | 0.020550 ± 0.001565 |
| | | | *S* | | |
| | | | *X* | | |

Table S2. The IUPAC names and chemical classes of the compounds in the dataset. Kovats indices were calculated using a series of *n*-alkanes (see *Materials and Methods*). Published KI values in plain text are from Yanez et al. 2002 in *Molecules*, doi: 10.3390/70900712. Published KI values in italics are from Chung et al. 1993 in *Journal of Agriculture and Food Chemistry*, doi: 10.1021/jf00034a033. When authentic standards were not available, we indicate the percentage match with the library for the compound, and list the top ten most abundant mass spectra (with their relative abundance in parentheses).

| Class | Class notes | Present in dynamic headspace | cylindrica headspace samples | unguiculata headspace samples | Present in SPME | Compound name | IUPAC | RT, min. | KI | Published KI | Standard run | Percent match with library | Mass spectra for tentatively identified compounds and unknowns: |
|---|---|---|---|---|---|---|---|---|---|---|---|---|---|
| Aromatics/Nitrogen | | x | 1 | 82 | x | 2-amino phenyl | 1-(2-aminophenyl)ethanone | 14.399 | 2265 | | Y | | |
| | | x | 4 | 14 | | 2-phenylethanol | 2-Phenylethan-1-ol | 12.268 | 1936 | *1873* | N | 97% | 50 (7.0), 51 (12.1), 63 (8.5), 65 (26.4), 77 (6.3), 78 (6.2), 91 (100), 93 |
| | Ester | x | 24 | 39 | | benzyl acetate | benzyl acetate | 10.911 | 1755 | | N | 91% | 43 (81.7), 51 (27.3), 65 (25.4), 77 (32.5), 79 (41.9), 89 (21.5), 90 (24.3), 91 (75.8), 108 (100), 150 (24.9) |
| | | x | 67 | 32 | x | benzyl alcohol | phenylmethanol | 12.001 | 1910 | *1837* | Y | | |
| | | x | 17 | 0 | | cinnamic alcohol | (E)-3-phenylprop-2-en-1-ol | 14.762 | 2311 | | N | 91% | 51 (55.8), 55 (37.8), 77 (49.0), 78 (71.2), 79 (37.3), 91 (84.0), 92 (100), 105 (47.2), 115 (49.3), 134 (48.7) |
| | methyl ester | x | 2 | 51 | | methyl nicotinate | methyl pyridine-3-carboxylate | 11.317 | 1737 | | Y | | |
| | organic ester | x | 8 | 25 | x | methyl salicylate | methyl 2-hydroxybenzoate | 11.320 | 1733 | | Y | | |
| | | Internal standard | 135 | 135 | | toluene | Methyl benzene | 4.184 | 1051 | *1042* | Y | | |
| | | x | 70 | 2 | | trans cinnamic | (E)-3-phenylprop-2-enal | 13.212 | 2092 | | Y | | |
| | Dimethyl ether | x | 86 | 0 | x | veratrole | 1,2-dimethoxybenzene | 10.974 | 1750 | | Y | | |
| Green Leafy Volatile | | x | 36 | 51 | x | 1 hexanol | Hexan-1-ol | 7.526 | 1374 | *1325* | Y | | |
| | | x | 22 | 71 | x | cis 3 hexen 1 ol | (Z)-hex-3-en-1-ol | 7.842 | 1404 | *1357* | Y | | |
| | | x | 15 | 88 | x | cis 3 hexenyl acetate | [(Z)-hex-3-enyl] acetate | 7.203 | 1335 | | Y | | |
| | | x | 14 | 17 | | cis jasmone | 3-methyl-2-[(Z)-pent-2-enyl]cyclopent-2-en-1-one | 12.583 | 1980 | | N | 92% | 41 (94.0), 53 (47.1), 67 (43.2), 55 (73.4), 77 (55.0), 79 (100), 91 (57.5), 93 (49.7), 110 (53.7), 122 (42.1) |
| | | x | 64 | 44 | x | trans 2 hexen 1 ol | (E)-hex-2-en-1-ol | 8.041 | 1431 | *1377* | Y | | |

(Table S2 continues)

#### Table S2 (continued)

| Class | Class notes | Present in dynamic headspace | cylindrica headspace samples | unguiculata headspace samples | Present in SPME | Compound name | IUPAC | RT, min. | KI | Published KI | Standard run | Percent match with library | Mass spectra for tentatively identified compounds and unknowns: |
|---|---|---|---|---|---|---|---|---|---|---|---|---|---|
| Monoterpenoids | unsaturated monoterpene | x | 16 | 49 | x | 2,6-dimethyl-1,3,5,7 octatetraene (trans) | (3E,5E)-2,6-dimethylocta-1,3,5,7-tetraene | 8.457 | 1466 | | N | <90% (low 80s) | 41 (51.2), 55 (28.2), 57 (100), 65 (20.7), 77 (38.6), 79 (31.4), 91 (96.8), 105 (19.5), 119 (63.4), 134 (35.0) |
| | irregular terpenoid | x | 41 | 36 | x | 6-methyl-5-hepten-2-one | 6-methylhept-5-en-2-one | 7.408 | 1356 | *1319* | N | 96% | 41 (51.2), 43 (100), 53 (7.6), 55 (32.2), 58 (14.7), 69 (21.9), 93 (7.6), 108 (18.9), 111 (7.4), 126 (2.0) |
| | | x | 8 | 68 | x | alpha pinene | 4,6,6-trimethylbicyclo[3.1.1]hept-3-ene | 4.009 | 1013 | 1092; *1034* | Y | | |
| | | x | 20 | 16 | x | alpha terpinene | 1-methyl-4-propan-2-ylcyclohexa-1,3-diene | 5.296 | 1155 | *1178* | N | 93% | 41 (43.2), 43 (19.4), 53 (19.9), 77 (51.6), 79 (50.2), 91 (56.6), 93 (100), 105 (23.0), 121 (70.6), 136 (33.6) |
| | Terpene alcohol | x | 27 | 30 | x | alpha terpineol | 2-(4-methylcyclohex-3-en-1-yl)propan-2-ol | 10.648 | 1722 | 1661; *1669* | Y | | |
| | | x | 3 | 10 | x | alpha terpinolene | 1-methyl-4-propan-2-ylidenecyclohexene | 6.865 | 1302 | *1275* | N | <90% | 45 (56.3), 51 (8.8), 58 (75.8), 77 (43.5), 79 (45.3), 91 (53.0), 93 (100), 105 (19.1), 121 (60.7), 136 (49.1) |
| | | x | 43 | 86 | x | beta myrcene | 7-methyl-3-methylideneocta-1,6-diene | 5.581 | 1169 | 1166; *1161* | Y | | |
| | | x | 47 | 44 | x | beta phellandrene | 3-methylidene-6-propan-2-ylcyclohexene | 6.065 | 1227 | *1209* | N | 95% | 41 (20.7), 65 (11.4), 77 (43.2), 79 (24.9), 80 (11.5), 91 (47.2), 92 (11.6), 93 (100), 94 (14.0), 136 (15.6) |
| | | x | 24 | 68 | x | beta pinene | 6,6-dimethyl-4-methylidenebicyclo[3.1.1]heptane | 4.942 | 1111 | 1136; *1114* | Y | | |
| | Terpene alcohol | x | 9 | 66 | | borneol | 4,7,7-trimethylbicyclo[2.2.1]heptan-3-ol | 9.468 | 1579 | *1677* | N | <90% | 45 (30.8), 53 (28.6), 67 (31.8), 77 (53.2), 79 (52.7), 81 (31.5), 91 (68.1), 93 (70.0), 95 (100), 150 (38.6) |
| | | x | 64 | 96 | x | cis beta ocimene | (3Z)-3,7-dimethylocta-1,3,6-triene | 6.346 | 1253 | *1225* | Y | | |
| | | x | 19 | 39 | x | gamma terpinene | 1-methyl-4-propan-2-ylcyclohexa-1,4-diene | 6.474 | 1261 | *1238* | Y | | |
| | Terpene alcohol | x | 43 | 14 | | geraniol | (2E)-3,7-dimethylocta-2,6-dien-1-ol | 10.492 | 1702 | *1814* | N | 90% | 41 (100), 53 (10.1), 59 (8.6), 67 (14.9), 68 (18.6), 69 (76.9), 81 (7.2), 93 (5.9), 111 (17.8), 123 (6.8) |
| | | x | 35 | 45 | x | limonene | 1-Methyl-4-(prop-1-en-2-yl)cyclohex-1-ene | 5.951 | 1300 | 1217 | Y | | |
| | Terpene alcohol | x | 14 | 21 | x | linalool | 3,7-Dimethylocta-1,6-dien-3-ol | 9.338 | 1563 | 1517; *1522* | Y | | |
| | Oxygenated | x | 26 | 33 | x | myroxide | 2,2-dimethyl-3-[(2E)-3-methylpenta-2,4-dienyl]oxirane | 8.831 | 1506 | | N | <90% (low 80s) | 41 (22.4), 43 (53.3), 53 (13.4), 56 (40.0), 73 (18.2), 77 (27.7), 79 (100), 81 (38.3), 84 (12.9), 93 (11.1) |
| | monoterpene aromatic | x | 0 | 23 | x | para cymene | 1-methyl-4-propan-2-ylbenzene | 6.725 | 1282 | *1261* | Y | | |
| | | x | 26 | 52 | x | sabinene | 4-methylidene-1-propan-2-ylbicyclo[3.1.0]hexane | 5.103 | 1116 | *1123* | Y | | |
| | Terpene alcohol | x | 1 | 22 | x | sabinene hydrate | 4-methyl-1-propan-2-ylbicyclo[3.1.0]hexan-4-ol | 9.400 | 1571 | | N | 90% | 43 (100), 45 (14.7), 53 (11.8), 69 (33.6), 71 (41.7), 79 (22.8), 81 (31.1), 93 (28.0), 111 (19.6), 121 (13.2) |
| | Terpene alcohol | x | 0 | 10 | x | terpinen-4-ol | 4-methyl-1-propan-2-ylcyclohex-3-en-1-ol | 9.883 | 1628 | *1579* | Y | | |
| | | x | 67 | 103 | x | trans beta ocimene | (3E)-3,7-dimethylocta-1,3,6-triene | 6.529 | 1274 | 1250; *1242* | Y | | |
| | Oxygenated | x | 0 | 4 | x | verbenone | 2,6,6-trimethylbicyclo[3.1.1]hept-2-en-4-one | 10.804 | 1741 | | N | 90% | 41 (67.1), 53 (39.0), 55 (36.8), 67 (42.9), 77 (38.0), 79 (53.0), 80 (61.6), 91 (77.1), 107 (100), 135 (51.6) |

(Table S2 continues)

### Table S2 (continued)

| Class | Class notes | Present in dynamic headspace | cylindrica headspace samples | unguiculata headspace samples | Present in SPME | Compound name | IUPAC | RT, min. | KI | Published KI | Standard run | Percent match with library | Mass spectra for tentatively identified compounds and unknowns: |
|---|---|---|---|---|---|---|---|---|---|---|---|---|---|
| Sesquiterpenes and C15 derivatives | | x | 47 | 22 | x | alloaromadendrene | (4aS,7R,7aR)-1,1,7-trimethyl-4-methylidene-2,3,4a,5,6,7,7a,7b-octahydro-1aH-cyclopropa[e]azulene | 10.365 | 1669 | *1639* | Y | | |
| | | x | 46 | 15 | x | alpha bergamotene | 4,6-dimethyl-6-(4-methylpent-3-enyl)bicyclo[3.1.1]hept-3-ene | 9.783 | 1616 | | N | 93% | 41 (72.4), 55 (43.2), 69 (36.8), 77 (35.9), 79 (35.2), 91 (44.5), 93 (100), 105 (25.5), 107 (30.4), 119 (83.8) |
| | | x | 65 | 42 | x | alpha humulene | (1E,4E,8E)-2,6,6,9-tetramethylcycloundeca-1,4,8-triene | 10.528 | 1699 | 1641; *1663* | Y | | |
| | | x | 132 | 14 | x | beta cadinene | (1S,4aR,8aS)-4,7-dimethyl-1-propan-2-yl-1,2,4a,5,8,8a-hexahydronaphthalene | 10.693 | 1713 | | Y | | |
| | | x | 9 | 6 | | beta farnesene | (6E)-7,11-dimethyl-3-methylidenedodeca-1,6,10-triene | 10.352 | 1678 | *1658* | Y | | |
| | | x | 85 | 34 | x | beta longipinene | (1S,2R,7S,8S)-2,6,6-Trimethyl-9-methylenetricyclo[5.4.0.02,8]undecane | 10.925 | 1757 | | N | <90% | 41 (100), 67 (36.1), 79 (69.2), 91 (65.9), 93 (59.4), 105 (54.12), 107 (55.4), 119 (36.8), 133 (37.2), 161 (64.8) |
| | Oxygenated | x | 38 | 7 | | caryophyllene oxide | (1*R*,4*R*,6*R*,10*S*)-9-Methylene-4,12,12-trimethyl-5-oxatricyclo[8.2.0.$0^{4,6}$]dodecane | 12.928 | 2029 | *1962* | N | <90% (low 80s) | 41 (100), 67 (37.2), 77 (45.6), 79 (77.7), 91 (53.0), 93 (66.5), 95 (43.6), 96 (36.2), 105 (40.0), 107 (33.4) |
| | | x | 16 | 9 | x | cis DMNT | (3Z)-4,8-dimethylnona-1,3,7-triene | 6.777 | 1293 | | N | 93% | 41 (100), 53 (14.9), 67 (8.9), 69 (96.7), 79 (12.5), 81 (9.7), 82 (6.5), 94 (4.3), 107 (7.7), 135 (2.6) |
| | | x | 4 | 39 | x | gamma cadinene | (1S,4aR,8aR)-7-methyl-4-methylidene-1-propan-2-yl-2,3,4a,5,6,8a-hexahydro-1H-naphthalene | 11.199 | 1791 | 1759; *1752* | N | <90% | 41 (81.5), 55 (36.28), 81 (48.7), 91 (63.6), 105 (83.5), 119 (90.9), 134 (90.4), 161 (100), 189 (26.1), 204 (33.0) |
| | | x | 59 | 67 | x | germacrene D | (1E,6E)-1-methyl-5-methylidene-8-propan-2-ylcyclodeca-1,6-diene | 10.885 | 1737 | 1687; 1705 | Y | | |
| | Oxygenated | x | 126 | 10 | x | intermedeol | (1S,4aS,7R,8aS)-1,4a-dimethyl-7-prop-1-en-2-yl-2,3,4,5,6,7,8,8a-octahydronaphthalen-1-ol | 14.560 | 2278 | | N | 95% | 41 (52.8), 43 (100), 55 (29.8), 67 (32.7), 71 (29.0), 81 (37.9), 93 (19.8), 161 (19.9), 189 (19.5), 204 (19.1) |
| | | x | 6 | 1 | | patchoulane | 1H-3a,7-Methanoazulene, octahydro-1,4,9,9-tetramethyl- | 13.994 | 2189 | | N | <90% (low 80s) | 41 (100), 53 (42.8), 55 (54.8), 67 (45.8), 79 (44.4), 91 (50.5), 93 (38.1), 95 (38.2), 105 (39.1), 119 (32.7) |
| | | x | 63 | 73 | x | trans beta caryophyllene | (1R,4E,9S)-4,11,11-trimethyl-8-methylidenebicyclo[7.2.0]undec-4-ene | 9.920 | 1626 | 1575; *1594* | Y | | |
| | | x | 1 | 20 | | trans DMNT | (3E)-4,8-dimethylnona-1,3,7-triene | 7.106 | 1319 | | Y | | |
| | | x | 26 | 53 | x | trans trans alpha farnesene | (3E,6E)-3,7,11-trimethyldodeca-1,3,6,10-tetraene | 11.068 | 1762 | *1725* | Y | | |
| | | x | 1 | 12 | x | unknown C15H24 (1) | | 10.671 | 1724 | | N | NA | 43 (86.0), 57 (100), 71 (70.6), 77 (13.7), 79 (12.9), 85 (27.4), 93 (27.8), 99 (12.0), 105 (19.6), 161 (26.2) |
| | | x | 66 | 4 | x | unknown C15H24 (2) | | 10.927 | 1757 | | N | NA | 41 (100), 77 (43.8), 79 (61.1), 91 (66.2), 93 (57.3), 105 (57.9), 107 (47.2), 119 (40.2), 133 (31.3), 161 (57.5) |
| | | x | 113 | 15 | x | unknown C15H24 (3) | | 11.283 | 1799 | | N | | 41 (81.2), 55 (44.0), 79 (38.2), 81 (37.0), 91 (48.1), 93 (30.6), 105 (38.6), 107 (75.8), 122 (88.0), 161 (100) |

Table S3. The 54 compounds in the dataset and their correlations with capscale axes 1 and 2. The *P* values (corrected for multiple tests, see *Materials and Methods*) are given for each correlation, as well as a column that lists the significance of each correlation at α = 0.01. The Trans. column give the type of transformation that was applied for subsequent univariate analyses: log = log-transformation, sqrt = square root-transformation, none = no transformation.

| **Compound** | **Cor. with cap1** | ***P* cap1** | **Sig. cap1** | **Cor. with cap2** | ***P* cap2** | **Sig. cap2** | **Trans.** |
|---|---|---|---|---|---|---|---|
| 1 hexanol | 0.1045 | 1.38E-01 | No | -0.0613 | 4.86E-01 | No | NA |
| 2 dimethyl 1 3 5 7 octatetraene trans | 0.3590 | 7.39E-09 | Yes | 0.3291 | 4.12E-07 | Yes | log |
| 2 amino phenyl ethanone | 0.4807 | 4.60E-16 | Yes | 0.2602 | 1.01E-04 | Yes | sqrt |
| 2 phenyl ethanol | 0.1233 | 7.99E-02 | No | -0.0370 | 6.85E-01 | No | NA |
| 6 methyl 5 hepten 2 one | -0.1424 | 4.15E-02 | No | -0.0624 | 4.86E-01 | No | NA |
| alloaromadendrene | 0.0035 | 9.55E-01 | No | 0.1265 | 1.20E-01 | No | NA |
| α-bergamotene | -0.1897 | 4.71E-03 | Yes | -0.0012 | 9.99E-01 | No | log |
| α-humulene | -0.0561 | 4.40E-01 | No | -0.0138 | 9.24E-01 | No | NA |
| α-pinene | 0.2016 | 2.75E-03 | Yes | -0.2518 | 1.54E-04 | Yes | log |
| α-terpinene | 0.0284 | 6.93E-01 | No | -0.1153 | 1.37E-01 | No | NA |
| α-terpineol | 0.1121 | 1.11E-01 | No | -0.0618 | 4.86E-01 | No | NA |
| α-terpinolene | 0.1032 | 1.40E-01 | No | -0.1162 | 1.37E-01 | No | NA |
| benzyl acetate | 0.0628 | 3.82E-01 | No | 0.0094 | 9.48E-01 | No | NA |
| benzyl alcohol | -0.1679 | 1.46E-02 | No | -0.0469 | 6.29E-01 | No | NA |
| β-cadinene | -0.6415 | 2.87E-31 | Yes | 0.0223 | 8.39E-01 | No | none |
| β-farnesene | 0.0893 | 2.12E-01 | No | 0.0432 | 6.38E-01 | No | NA |
| β-longipinene | -0.3758 | 1.18E-09 | Yes | 0.0307 | 7.52E-01 | No | sqrt |
| β-myrcene | 0.2861 | 6.79E-06 | Yes | -0.0297 | 7.52E-01 | No | sqrt |
| β-phellandrene | 0.0344 | 6.32E-01 | No | -0.1224 | 1.20E-01 | No | NA |
| β-pinene | 0.1980 | 3.22E-03 | Yes | -0.2580 | 1.06E-04 | Yes | log |
| borneol | 0.3563 | 9.02E-09 | Yes | 0.3131 | 1.62E-06 | Yes | sqrt |
| caryophyllene oxide | -0.0398 | 5.79E-01 | No | 0.0447 | 6.38E-01 | No | NA |

(Table S3 continues)

Table S3 (continued)

| **Compound** | **Cor. with cap1** | ***P* cap1** | **Sig. cap1** | **Cor. with cap2** | ***P* cap2** | **Sig. cap2** | **Trans.** |
|---|---|---|---|---|---|---|---|
| cinnamic alcohol | -0.0780 | 2.71E-01 | No | 0.0626 | 4.86E-01 | No | NA |
| *Z*-3-hexen-1-ol | 0.1571 | 2.38E-02 | No | -0.2750 | 3.52E-05 | Yes | log |
| *Z*-3-hexenyl acetate | 0.3767 | 1.18E-09 | Yes | -0.3495 | 6.44E-08 | Yes | log |
| *Z*-β-ocimene | 0.5282 | 1.51E-19 | Yes | 0.4436 | 5.16E-13 | Yes | sqrt |
| *Z*-jasmone | 0.0532 | 4.61E-01 | No | -0.1381 | 7.85E-02 | No | NA |
| *Z*-dimethylnonatriene | -0.0414 | 5.72E-01 | No | 9.23E-05 | 9.99E-01 | No | NA |
| γ-cadinene | 0.2280 | 5.30E-04 | Yes | 0.0792 | 4.04E-01 | No | sqrt |
| γ-terpinene | 0.1917 | 4.42E-03 | Yes | -0.2311 | 6.27E-04 | Yes | log |
| geraniol | -0.0717 | 3.09E-01 | No | 0.0739 | 4.52E-01 | No | NA |
| germacrene D | 0.0887 | 2.12E-01 | No | -0.0691 | 4.81E-01 | No | NA |
| intermedeol | -0.6455 | 1.73E-31 | Yes | -0.0148 | 9.24E-01 | No | sqrt |
| limonene | 0.0222 | 7.44E-01 | No | -0.1235 | 1.20E-01 | No | NA |
| linalool | 0.0772 | 2.71E-01 | No | -0.1249 | 1.20E-01 | No | NA |
| methyl nicotinate | 0.2953 | 3.27E-06 | Yes | 0.1546 | 3.95E-02 | No | log |
| methyl salicylate | 0.0427 | 5.69E-01 | No | 0.1030 | 1.97E-01 | No | NA |
| myroxide | 0.1263 | 7.33E-02 | No | 0.2285 | 6.83E-04 | Yes | log |
| para cymene | 0.1540 | 2.65E-02 | No | -0.1089 | 1.67E-01 | No | NA |
| patchoulane | -0.0880 | 2.12E-01 | No | -0.0426 | 6.38E-01 | No | NA |
| sabinene | 0.1382 | 4.80E-02 | No | -0.1823 | 1.10E-02 | No | NA |
| sabinene hydrate | 0.1089 | 1.21E-01 | No | -0.2749 | 3.52E-05 | Yes | log |
| terpinen-4-ol | 0.1183 | 9.16E-02 | No | -0.1689 | 2.07E-02 | No | NA |
| *E*-2-hexen-1-ol | -0.0839 | 2.34E-01 | No | -0.0486 | 6.22E-01 | No | NA |
| *E*-β-caryophyllene | -0.0259 | 7.11E-01 | No | -0.0531 | 5.77E-01 | No | NA |
| *E*-β-ocimene | 0.5261 | 1.70E-19 | Yes | 0.4573 | 1.26E-13 | Yes | sqrt |
| *E*-cinnamic aldehyde | -0.2952 | 3.27E-06 | Yes | 0.0416 | 6.38E-01 | No | sqrt |

(Table S3 continues)

Table S3 (continued)

| **Compound** | **Cor. with cap1** | ***P* cap1** | **Sig. cap1** | **Cor. with cap2** | ***P* cap2** | **Sig. cap2** | **Trans.** |
|---|---|---|---|---|---|---|---|
| *E*-dimethylnonatriene | 0.1181 | 9.16E-02 | No | 0.0675 | 4.85E-01 | No | NA |
| *E*-*E*-α-farnesene | 0.1436 | 4.11E-02 | No | 0.0029 | 9.99E-01 | No | NA |
| unknown $C_{15}H_{24}$ #1 | 0.1316 | 6.14E-02 | No | -0.0058 | 9.79E-01 | No | NA |
| unknown $C_{15}H_{24}$ #2 | -0.3326 | 1.05E-07 | Yes | 0.0706 | 4.78E-01 | No | sqrt |
| unknown $C_{15}H_{24}$ #3 | -0.5157 | 1.04E-18 | Yes | 0.0122 | 9.27E-01 | No | sqrt |
| veratrole | -0.2723 | 2.02E-05 | Yes | 0.0636 | 4.86E-01 | No | sqrt |
| verbenone | -0.0122 | 8.58E-01 | No | -0.1175 | 1.37E-01 | No | NA |

Table S4. The values for the centroids for each species and community type along CAP axes 1 and 2.

| **Species & Community Type** | **CAP axis 1** | **CAP axis 2** |
|---|---|---|
| *C. cylindrica* | | |
| Single-species communities | -1.006 | 0.137 |
| Two-species communities | -0.678 | 0.196 |
| Four-species communities | -0.547 | -0.391 |
| *C. unguiculata* | | |
| Single-species communities | 0.676 | -1.212 |
| Two-species communities | 0.857 | -0.061 |
| Four-species communities | 0.698 | 1.330 |

Table S5. Tukey's Honest Significant Difference post-hoc tests on the differences between *C. cylindrica* and *C. unguiculata* in the given compound or compound class at each community type. The type of data transformation (log or square root) is indicated in the trait column. Compounds and compound classes are abbreviated as follows: SES: sesquiterpenoids; GLV: "green leafy volatiles;" AROM: aromatics; 2-APE: 2-amino phenyl ethanone; α-P: α-pinene; β-P: β-pinene; γ-T: γ-terpinene; SH: sabinene hydrate; MN: methyl nicotinate; (*E*)-C A: (*E*)-cinnamic aldehyde; V: veratrole; (*Z*)-3-H A: (*Z*)-3-hexenyl acetate. For log-transformed traits, tests are performed on the log odds scale such that ratios greater than one indicate that *C. cylindrica* has a higher emission rate than *C. unguiculata*, and ratios lower than one indicate that *C. unguiculata* has a higher emission rate. For square root-transformed traits, estimates are back-transformed from the square-root scale such that positive estimates indicate that *C. cylindrica* has a higher emission rate than *C. unguiculata*. General linear hypothesis tests determine the differences between the differences at a pair of community types. Positive estimates indicate the first community type in the hypothesis has a larger difference in the trait relative to the second community type in the hypothesis. All tests were corrected for multiple comparisons.

| **Tukey's Honest Significant Difference tests** *Testing differences between species at com. types* | | | | | **General linear hypothesis tests** *Testing differences of the differences* | | | |
|---|---|---|---|---|---|---|---|---|
| Trait | Com. Type | Estimate or ratio ± 1 SE | *t* ratio | *P* | Hypo. | Estimate ± 1 SE | Z value | *P* |
| log SES | One | 57.430 ± 21.44 | 10.85 | **<0.001** | Δ One = Δ Two | 2.1868 ± 0.5229 | 4.18 | **< 0.001** |
| | Two | 6.450 ± 2.36 | 5.090 | **<0.001** | Δ One = Δ Four | 1.0727 ± 0.5229 | 2.05 | 0.1001 |
| | Four | 19.650 ± 7.19 | 8.132 | **<0.001** | Δ Two = Δ Four | -1.1141 ± 0.5179 | -2.15 | 0.0797 |
| log GLV | One | 0.070 ± 0.043 | -4.269 | **0.001** | Δ One = Δ Two | 1.043 ± 0.802 | 1.30 | 0.3935 |
| | Two | 0.025 ± 0.012 | -7.370 | < **0.001** | Δ One = Δ Four | -3.029 ± 0.802 | -3.78 | **<0.001** |
| | Four | 1.439 ± 0.724 | 0.723 | 0.4704 | Δ Two = Δ Four | -4.073 ± 0.712 | -5.72 | < **0.001** |
| sqrt AROM | One | -0.004 ± 0.011 | -0.374 | 0.713 | Δ One = Δ Two | -0.041 ± 0.015 | 2.760 | **0.016** |
| | Two | 0.036 ± 0.010 | 3.662 | **<0.001** | Δ One = Δ Four | -0.005 ± 0.015 | -0.35 | 0.934 |
| | Four | 0.001 ± 0.010 | 0.115 | 0.909 | Δ Two = Δ Four | 0.035 ± 0.014 | -2.51 | **0.032** |
| sqrt 2-APE | One | -0.009 ± 0.005 | -1.855 | 0.0954 | Δ One = Δ Two | 0.017 ± 0.006 | 2.918 | 0.009 |
| | Two | -0.026 ± 0.003 | -9.870 | < **0.001** | Δ One = Δ Four | 0.021 ± 0.006 | 3.731 | **<0.001** |

(Table S5 continues)

Table S5 (continued)

| Trait | Com. Type | Estimate or ratio ± 1 SE | *t* ratio | *P* | H | Estimate ± 1 SE | Z value | *P* |
|---|---|---|---|---|---|---|---|---|
| sqrt 2-APE | Four | -0.031 ± 0.003 | -11.63 | < **0.001** | Δ Two = Δ Four | 0.005 ± 0.004 | 1.243 | 0.421 |
| log α-P | One | 0.213 ± 0.061 | -5.406 | < **0.001** | Δ One = Δ Two | -0.347 ± 0.341 | -1.02 | 0.562 |
| | Two | 0.301 ± 0.056 | -6.514 | < **0.001** | Δ One = Δ Four | -1.002 ± 0.341 | -2.94 | **0.009** |
| | Four | 0.580 ± 0.107 | -2.958 | **0.0034** | Δ Two = Δ Four | -0.656 ± 0.261 | -2.51 | **0.031** |
| log β-P | One | 0.627 ± 0.051 | -5.765 | < **0.001** | Δ One = Δ Two | -0.105 ± 0.107 | -0.97 | 0.595 |
| | Two | 0.696 ± 0.049 | -5.150 | < **0.001** | Δ One = Δ Four | -0.358 ± 0.107 | -3.34 | **0.002** |
| | Four | 0.897 ± 0.063 | -1.538 | 0.125 | Δ Two = Δ Four | -0.254 ± 0.099 | -2.55 | **0.029** |
| log γ-T | One | 0.579 ± 0.072 | -4.390 | < **0.001** | Δ One = Δ Two | 0.044 ± 0.176 | 0.248 | 0.967 |
| | Two | 0.554 ± 0.069 | -4.742 | < **0.001** | Δ One = Δ Four | -0.495 ± 0.176 | -2.81 | **0.014** |
| | Four | 0.949 ± 0.118 | -0.420 | 0.675 | Δ Two = Δ Four | -0.538 ± 0.176 | -3.06 | **0.006** |
| log SH | One | 0.759 ± 0.055 | -3.821 | **0.002** | Δ One = Δ Two | -0.029 ± 0.096 | -0.31 | 0.950 |
| | Two | 0.782 ± 0.050 | -3.880 | < **0.001** | Δ One = Δ Four | -0.276 ± 0.096 | -2.87 | **0.011** |
| | Four | 1.000 ± 0.064 | 0 | 1 | Δ Two = Δ Four | -0.246 ± 0.090 | -2.74 | **0.017** |
| log MN | One | 0.783 ± 0.146 | -1.313 | 0.216 | Δ One = Δ Two | 0.684 ± 0.228 | 2.999 | **0.008** |
| | Two | 0.395 ± 0.051 | -7.086 | < **0.001** | Δ One = Δ Four | 0.494 ± 0.228 | 2.168 | 0.076 |
| | Four | 0.477 ± 0.063 | -5.639 | < **0.001** | Δ Two = Δ Four | -0.190 ± 0.185 | -1.02 | 0.5603 |
| sqrt (*E*)-C A | One | 0.008 ± 0.003 | 2.525 | 0.021 | Δ One = Δ Two | -0.013 ± 0.004 | -2.95 | **0.009** |
| | Two | 0.021 ± 0.003 | 6.859 | < **0.001** | Δ One = Δ Four | -0.009 ± 0.004 | -2.04 | 0.1033 |
| | Four | 0.016 ± 0.003 | 5.540 | < **0.001** | Δ Two = Δ Four | 0.004 ± 0.004 | 0.932 | 0.6199 |

(Table S5 continues)

Table S5 (continued)

| Trait | Com. Type | Estimate or ratio ± 1 SE | *t* ratio | *P* | | H | Estimate ± 1 SE | Z value | *P* |
|---|---|---|---|---|---|---|---|---|---|
| sqrt V | One | 0.007 ± 0.003 | 2.175 | 0.042 | | Δ One = Δ Two | -0.041± 0.004 | -9.26 | < **0.001** |
| | Two | 0.047 ± 0.003 | 15.28 | < **0.001** | | Δ One = Δ Four | -0.023 ± 0.004 | -5.22 | < **0.001** |
| | Four | 0.030 ± 0.003 | 9.567 | < **0.001** | | Δ Two = Δ Four | 0.018 ± 0.004 | 4.036 | < **0.001** |
| log (*Z*)-3-H A | One | 0.015 ± 0.009 | -7.054 | < **0.001** | | Δ One = Δ Two | -0.198 ± 0.697 | -0.28 | 0.956 |
| | Two | 0.019 ± 0.007 | -10.80 | < **0.001** | | Δ One = Δ Four | -4.520 ± 0.697 | -6.48 | < **0.001** |
| | Four | 1.411 ± 0.520 | 0.935 | 0.351 | | Δ Two = Δ Four | -4.322 ± 0.521 | -8.30 | < **0.001** |

Table S6. Outputs of ANOVAs for the nine compounds that had significant community type x species interactions. The trait column indicates the type of data transformation applied to the compound. Compounds are abbreviated as follows: 2-APE: 2-amino phenyl ethanone; α-P: α-pinene; β-P: β-pinene; γ-T: γ-terpinene; SH: sabinene hydrate; MN: methyl nicotinate; (*E*)-C A: (*E*)-cinnamic aldehyde; V: veratrole; (*Z*)-3-H A: (*Z*)-3-hexenyl acetate. *P* values for the community type x species interactions are adjusted for performing 23 tests (e.g. univariate analyses on all compounds that were significantly correlated with one or both of the first two CAP axes).

| **Trait** | **Term** | **MS** | **NDF** | **DDF** | ***F*** | ***P*** |
|---|---|---|---|---|---|---|
| sqrt 2-APE | Type | 0.001 | 2 | 7.287 | 3.450 | 0.088 |
| | Species | 0.017 | 1 | 21.403 | 110.540 | 6.51 E-10 |
| | Type x Species | 0.001 | 2 | 20.725 | 6.960 | 0.022 |
| log α-P | Type | 1.709 | 2 | 6.219 | 2.236 | 0.186 |
| | Species | 55.313 | 1 | 29.081 | 72.358 | 2.21 E-09 |
| | Type x Species | 4.175 | 2 | 22.979 | 5.461 | 0.038 |
| log β-P | Type | 0.348 | 2 | 5.690 | 3.127 | 0.121 |
| | Species | 5.942 | 1 | 67.701 | 53.379 | 4.06 E-10 |
| | Type x Species | 0.693 | 2 | 40.299 | 6.225 | 0.022 |
| log γ-T | Type | 1.270 | 2 | 264 | 3.640 | 0.028 |
| | Species | 10.614 | 1 | 264 | 30.410 | 8.33 E-08 |
| | Type x Species | 2.011 | 2 | 264 | 5.761 | 0.022 |
| log SH | Type | 0.385 | 2 | 7.247 | 4.245 | 0.060 |
| | Species | 1.862 | 1 | 83.568 | 20.537 | 1.93 E-05 |
| | Type x Species | 0.487 | 2 | 51.073 | 5.374 | 0.029 |
| log MN | Type | 1.262 | 2 | 6.647 | 3.263 | 0.103 |
| | Species | 20.454 | 1 | 38.803 | 52.900 | 9.33E-09 |
| | Type x Species | 1.748 | 2 | 28.389 | 4.521 | 0.050 |
| sqrt (*E*)-C A | Type | 0.001 | 2 | 8.737 | 5.371 | 0.030 |
| | Species | 0.015 | 1 | 128.528 | 73.419 | 2.82E-14 |
| | Type x Species | 0.001 | 2 | 78.526 | 4.543 | 0.039 |
| sqrt V | Type | 0.009 | 2 | 264 | 43.138 | < 2.2E-16 |
| | Species | 0.053 | 1 | 264 | 243.318 | < 2.2E-16 |
| | Type x Species | 0.009 | 2 | 264 | 43.138 | 5.06 E-15 |
| log (*Z*)-3-H A | Type | 5.014 | 2 | 8.530 | 1.642 | 0.249 |
| | Species | 299.506 | 1 | 35.599 | 98.049 | 9.12 E-12 |
| | Type x Species | 125.399 | 2 | 29.843 | 41.052 | 3.12 E-08 |
